# Local adaptation and spatiotemporal patterns of genetic diversity revealed by repeated sampling of *Caenorhabditis elegans* across the Hawaiian Islands

**DOI:** 10.1101/2021.10.11.463952

**Authors:** Timothy A. Crombie, Paul Battlay, Robyn E. Tanny, Kathryn S. Evans, Claire M. Buchanan, Daniel E. Cook, Clayton M. Dilks, Loraina A. Stinson, Stefan Zdraljevic, Gaotian Zhang, Nicole M. Roberto, Daehan Lee, Michael Ailion, Kathryn A. Hodgins, Erik C. Andersen

## Abstract

The nematode *Caenorhabditis elegans* is among the most widely studied organisms, but relatively little is known about its natural ecology. Wild *C. elegans* have been isolated from both temperate and tropical climates, where they feed on bacteria associated with decomposing plant material. Genetic diversity is low across much of the globe but high in the Hawaiian Islands and across the Pacific Rim. The high genetic diversity found there suggests that: (1) the origin of the species lies in Hawaii or the surrounding Pacific Rim; and (2) the ancestral niche of the species is likely similar to the Hawaiian niche. A recent study of the Hawaiian niche found that genetically distinct groups appeared to correlate with elevation and temperature, but the study had a limited sample size. To better characterize the niche and genetic diversity of *C. elegans* on the Hawaiian Islands and to explore how genetic diversity might be influenced by local adaptation, we repeatedly sampled nematodes over a three-year period, measured various environmental parameters at each sampling site, and whole-genome sequenced the *C. elegans* isolates that we identified. We found that the typical Hawaiian *C. elegans* niche is moderately moist native forests at high elevations (500 to 1500 meters) where temperatures are cool (15 to 20°C). We measured levels of genetic diversity and differentiation among Hawaiian strains and found evidence of seven genetically distinct groups distributed across the islands. Then, we scanned these genomes for signatures of local adaptation and identified 18 distinct regions that overlap with hyperdivergent regions, which are likely maintained by balancing selection and enriched for genes related to environmental sensing, xenobiotic detoxification, and pathogen resistance. These results provide strong evidence of local adaptation among Hawaiian *C. elegans* and a possible genetic basis for this adaptation.

## Introduction

The nematode *Caenorhabditis elegans* is a powerful model organism that has facilitated advances in the fields of developmental, cellular, and molecular biology. Yet, despite decades of research, only recent attention has been given to its ecology, natural diversity, and evolutionary history (Frézal & Félix, 2015; Petersen, Dirksen, & Schulenburg, 2015; Schulenburg & Félix, 2017). Early surveys of wild *C. elegans* were largely focused on sampling from compost heaps in rural gardens. More recently, natural substrates in less artificial habitats have been surveyed revealing that wild *C. elegans* are typically found on rotting fruit and vegetable matter, where they persist by feeding on the diverse bacterial communities associated with these substrates (Schulenburg & Félix, 2017). However, intensive surveys of wild *C. elegans* in less artificial habitats have mostly been focused in Europe and the western continental United States, and relatively few strains have been isolated from other regions of the world (Barrière & Félix, 2005; Crombie et al., 2019; Félix & Duveau, 2012; Frézal & Félix, 2015; K. C. Kiontke et al., 2011; Petersen, Dirksen, Prahl, Strathmann, & Schulenburg, 2014; Richaud, Zhang, Lee, Lee, & Félix, 2018; Sivasundar & Hey, 2005).

As the number of wild isolates has grown, analysis of the genetic diversity revealed a striking pattern of exceptionally low diversity in strains isolated from across much of the globe and higher diversity in strains isolated from the Hawaiian Islands and more generally across the Pacific Rim (Crombie et al., 2019; Lee et al., 2021). This pattern is thought to be explained at least in part by recent chromosome-scale selective sweeps that have purged diversity from the species, presumably while the *C. elegans* range expanded in association with humans (Andersen et al., 2012). Genetic diversity in a species is often elevated in populations near the geographic origin of the species (Haipeng Li & Stephan, 2006; Nielsen et al., 2017; Peter et al., 2018). Therefore, surveys of the Hawaiian Islands and the Pacific Rim are especially important for understanding the evolutionary origins of *C. elegans* (Andersen et al., 2012; Crombie et al., 2019; Lee et al., 2021). Similarly, characterization of the niche in high diversity regions is especially relevant for understanding ancient patterns of adaptive differentiation in *C. elegans*, as many other regions have only recently been colonized and harbor limited genetic diversity. A recent survey of Hawaiian nematodes determined that *C. elegans* can be found on at least five of the major islands at elevations ranging from 500 to 1500 meters and at temperatures ranging from 15 to 25°C (Crombie et al., 2019). Although that analysis was based on a relatively small sample size, we observed a correlation between environmental variables at sampling locations and the genetic groups to which individuals were assigned. These findings raise the possibility that local adaptation to heterogeneous environments could contribute to the exceptional genetic diversity observed among Hawaiian strains.

Local adaptation can occur in response to varying selection pressures imposed by spatially heterogeneous environments and can cause alleles to vary in frequency across the range of a species (Kawecki & Ebert, 2004). For this reason, alleles correlated with features of the environment are often interpreted as a signature of local adaptation (Booker, Yeaman, & Whitlock, 2021; Coop, Witonsky, Di Rienzo, & Pritchard, 2010). However, disentangling the signatures of local adaptation from patterns caused by neutral forces and/or demographic histories can be difficult (Rellstab, Gugerli, Eckert, Hancock, & Holderegger, 2015). For this reason, various genotype-environment association (GEA) methods have been developed to detect genomic signatures of local adaptation while accounting for the effects of other forces where possible (Hoban et al., 2016; Rellstab et al., 2015). In recent years, these tools have been applied to help detect the genetic basis and modes of adaptation to environmental variation for several species that are important to the ecology, evolution, and conservation biology fields (Hancock et al., 2011; Rellstab et al., 2015).

To better characterize the exceptional patterns of genetic diversity on the Hawaiian Islands and to explore how this diversity might be influenced by local adaptation, we repeatedly sampled nematodes over a three-year period, measured various environmental niche parameters at each sampling site, and whole-genome sequenced *C. elegans* isolates. We found that *C. elegans* environmental niche preferences differ substantially from the other selfing *Caenorhabditis* species (*C. briggsae* and *C. tropicalis*) on the Hawaiian Islands. We also observed considerable environmental variation within the *C. elegans* niche itself. We measured genetic diversity and differentiation using whole-genome sequences from 464 Hawaiian *C. elegans* strains we collected and 36 Hawaiian strains that were collected by our collaborators. We found that this sample of Hawaiian strains comprises 163 non-redundant genome-wide haplotypes, which we refer to as isotypes, representing an almost four-fold increase in sample-size relative to the most recent study of Hawaiian diversity (Crombie et al., 2019). Using principal components analysis (PCA), we found evidence of seven genetically distinct groups within the sample. Additionally, we identified overlapping regions of the genome that display signatures of local adaptation to various environmental variables, including elevation, temperature, and precipitation, using two GEA methods. These results further our understanding of the evolutionary forces shaping genetic diversity on the Hawaiian Islands and provide clues about the functional genetic variation relevant to local adaptation.

## Materials and Methods

### Sampling strategy

We sampled Hawaiian nematodes on six occasions from August 2017 to January 2020. These sampling projects varied in size, with four larger projects, comprising more than 500 samples in August 2017, October 2018, October 2019, and December 2019, and two smaller projects, comprising fewer than 100 samples in August 2019 and January 2020. For each project, we chose sampling locations based on accessibility to hiking trails and by proximity to where *Caenorhabditis* nematodes had been collected previously (Andersen et al., 2012; Cook et al., 2016; Crombie et al., 2019; Hahnel et al., 2018; Hodgkin & Doniach, 1997). In the four larger projects, we attempted to sample broadly across different habitats, which in Hawaii tend to vary along gradients of elevation and exposure to trade winds. At each sampling location, we opportunistically sampled substrates known to harbor *Caenorhabditis* nematodes, including rotting fruits, seeds, nuts, flowers, stems, mixed vegetal litter, compost, wood, soil, fungus, live arthropods, and molluscs (Crombie et al., 2019; Ferrari et al., 2017; Schulenburg & Félix, 2017). The mixed vegetal litter category describes substrates that contain detritus or dead organic material that forms a layer over the soil at most collection sites.

### Field sampling and environmental data collection

We collected samples from nature by transferring substrate material directly into a pre-barcoded collection bag as described previously (Crombie et al., 2019). To characterize the abiotic niche of *Caenorhabditis* nematodes, we collected data for environmental parameters at each sampling site, including the surface temperature of the sample using an infrared thermometer (Lasergrip 1080, Etekcity, Anaheim, CA) and the ambient temperature and humidity near the sample using a combined thermometer and hygrometer device (GM1362, GoerTek, Weifang, China). We used a mobile device and a geographical data-collection application, Fulcrum, to record the environmental parameter values, substrate GPS coordinates, *in situ* photographs of the substrate, and categorical descriptions of the substrate in a cloud database. We then exported the collection data from the Fulcrum database and processed it using the easyFulcrum (v1.0.0) R package to flag and correct anomalous data records (Di Bernardo, Crombie, Cook, & Andersen, 2021). For *C. elegans* positive samples, we used a hierarchical clustering approach to group samples within a 3 km distance. We used the *distm()* function from the geosphere (v1.5-10) R package to calculate a geodesic distance matrix from the sample locations and then clustered the samples within 3 km groups using the *hclust()* and *cutree()* functions from the stats (v3.6.3) package. We chose to cluster with the 3 km distance because it reduced the within cluster sum of squares when samples were grouped by island and largely recapitulated the distinct hiking trail or region where the samples were collected.

### GIS environmental data

To further characterize the environmental conditions near each sampling site, we used publicly available geographic information system (GIS) data and processed these data in R (v3.6.3) with the raster (v3.1-5) and sf (v0.9-5) packages (Hijmans, 2020; Pebesma, 2018; R Core Team, 2020). We used GIS maps produced for the assessment of evapotranspiration in the state of Hawaii at 250 meter resolution to assess various average annual climate parameter values for our sampling sites, including air temperature, surface temperature, available soil moisture, and leaf area index (LAI) (T. W. Giambelluca, Shuai, Barnes, & Alliss, 2014). LAI quantifies the amount of vegetation in a given area as the ratio of one-sided leaf area to ground area. We also used GIS data from the Rainfall Atlas of Hawaii to determine mean annual rainfall totals at approximately 250 meter resolution (Frazier, Giambelluca, Diaz, & Needham, 2016; Thomas W. Giambelluca et al., 2013) and two GIS maps generated for the Carbon Assessment of Hawaii (CAH) to determine the land cover and habitat status at our sampling sites at 30 meter resolution (Jacobi, Price, Fortini, Gon, & Berkowitz, 2017). We use the term land cover to describe characteristics of the plant communities within each 30 meter map unit and the term habitat status to describe the condition of those plant communities. The CAH land cover map uses a hierarchical classification system that allows the user to group the mapped units into different configurations, including 48 detailed plant community units, 27 generalized land cover units, 13 biome units, and seven major land cover units. The CAH habitat status map depicts the distribution of plant communities that are (1) dominated by native species, (2) mixed native and alien species, (3) heavily disturbed areas with few native species, and (4) areas with less than five percent vegetation cover. We renamed classifications from the CAH habitat status map to native, introduced, disturbed, and bare. Bare habitats contain little vegetation mostly due to recent lava flows and we did not include this habitat class in enrichment analyses because it was sampled infrequently (10 samples) and *Caenorhabditis* nematodes were never found there.

### Nematode isolation

Following sample collection, bagged and barcoded samples were shipped overnight from Hawaii to Northwestern University where the substrates were transferred from the barcoded collection bags to matching barcoded 10 cm NGMA plates (Andersen, Bloom, Gerke, & Kruglyak, 2014) seeded with OP50 bacteria. We attempted to isolate nematodes from these collection plates two days after the substrates were transferred. If no nematodes were found, we attempted to isolate nematodes again after seven days. For each collection plate, up to seven gravid nematodes were isolated by transferring them individually to pre-barcoded 3.5 cm NGMA isolation plates seeded with OP50 bacteria. In many cases, gravid adult animals were not found on the collection plates, so we isolated larval stages instead. This technique biases our isolation strategy towards selfing nematode species. At the time of isolation, we scanned the barcodes on the collection and isolation plates with the Fulcrum mobile app so that each isolate was linked to the appropriate field collection record in the Fulcrum database. If we could not find nematodes on the collection plate after seven days, we recorded that the isolated nematode failed to proliferate. We exported the isolation data from the Fulcrum database and processed it with the easyFulcrum R package to join the isolation records with the collection records for further analysis (Di Bernardo et al., 2021).

### Nematode Identification

We identified *Caenorhabditis* isolates to the species-level and isolates of other genera to the genus-level by analysis of the Internal Transcribed Spacer (ITS2) region between the 5.8S and 28S rDNA genes (Barrière & Félix, 2014; K. C. Kiontke et al., 2011). The isolated nematodes were stored at 20°C for up to 21 days before they were genotyped but were not passaged during this time to avoid multiple generations of proliferation. For genotyping, we lysed three to five nematodes from an isolation plate in 8 µL of lysis solution (100 mM KCl, 20 mM Tris pH 8.2, 5 mM MgCl2, 0.9% IGEPAL, 0.9% Tween 20, 0.02% gelatin with proteinase K added to a final concentration of 0.4 mg/ml) then froze the solution at -80°C for up to 12 hours. If isolated nematodes could not be found on the isolation plates, we categorized them as “Not genotyped”. We loaded 2 µL of thawed lysis material into 40 µL reactions with primers spanning a portion of the ITS2 region using forward primer oECA1687 (CTGCGTTATTTACCACGAATTGCARAC) and reverse primer oECA202 (GCGGTATTTGCTACTACCAYYAMGATCTGC) (K. C. Kiontke et al., 2011). We also loaded 2 µL of the lysed material into 40 µL reactions with a second set of primers that amplify about 500 bp of 18S rDNA in Rhabditid nematodes using forward primer oECA1271 (TACAATGGAAGGCAGCAGGC) and reverse primer oECA1272 (CCTCTGACTTTCGTTCTTGATTAA) (Haber et al., 2005). The PCR conditions for both primer sets were described previously (Crombie et al., 2019). Products from both PCR amplifications were visualized on a 2% agarose gel in 1X TAE buffer. We classified isolates that did not produce bands with the either primer set as “unknown nematode” and isolates for which the 18S region amplified, but the ITS2 region did not, as “non-*Caenorhabditis*”. The isolates that produced bands with both primer sets were investigated further using Sanger sequencing of the ITS2 PCR products with forward primer oECA306 (CACTTTCAAGCAACCCGAC). We compared these ITS2 sequences to the National Center for Biotechnology Information (NCBI) database using the BLASTn algorithm, which identified *Caenorhabditis* isolates to the species-level. Isolates with sequences that aligned best to genera other than *Caenorhabditis* were only identified to the genus-level. In most cases, isolates identified as *C. briggsae*, *C. elegans*, or *C. tropicalis* were named and cryopreserved. For each named strain, one of four recently starved 10 cm NGMA plates was used to cryopreserve the strain, and the other three plates were used for DNA extraction and whole-genome sequencing.

### Illumina library construction and whole-genome sequencing

To extract DNA, we transferred nematodes from three recently starved 10 cm NGMA plates originally spotted with OP50 *E. coli* into a 15 ml conical tube by washing with 10 mL of M9. We then used gravity to settle animals in the conical tube, removed the supernatant, and added 10 mL of fresh M9. We repeated this wash method three times to serially dilute the *E. coli* in the M9 and allow the animals time to purge ingested *E. coli*. Genomic DNA was isolated from 100 to 300 µl nematode pellets using the Blood and Tissue DNA isolation kit (cat# 69506, QIAGEN, Valencia, CA) following established protocols (Cook et al., 2016). The DNA concentration was determined for each sample using the Qubit dsDNA Broad Range Assay Kit (cat# Q32850, Invitrogen, Carlsbad, CA). Sequencing libraries were either generated with KAPA Hyper Prep kits (Kapa Biosystems, Wilmington, MA), Illumina Nextera Sample Prep Kit (Illumina, Inc. San Diego, CA), or New England BioLabs NEBNext® Ultra™ II FS DNA Library Prep (NEB, Ipswich, MA). Samples were sequenced at the Duke Center for Genomic and Computational Biology, Novogene, or the Northwestern Sequencing facility, NUSeq. All samples were sequenced on the Illumina HiSeq 4000 or NovaSeq 6000 platform (paired-end 150 bp reads). The raw sequencing reads for strains used in this project are available from the NCBI Sequence Read Archive (Project PRJNA549503).

### Variant calling

To ensure reproducible data analysis, all genomic analyses were performed using pipelines generated in the Nextflow workflow management system framework (Di Tommaso et al., 2017). Full descriptions of the pipelines can be found on the Andersen laboratory dry guide (http://andersenlab.org/dry-guide/latest/pipeline-overview/). Raw sequencing reads were trimmed using fastp, which removed low-quality bases and adapter sequences with the trim-fq-nf. The trimmed reads were mapped with BWA (v0.7.17) (Heng Li, 2013) to the N2 reference genome (WS276) with the alignment-nf. Next, we called SNVs using the wi-gatk pipeline with GATK with HaplotypeCaller and jointly called with GenotypeGVCFs. Because *C. elegans* is a selfing species and we expect the vast majority of the sites to be homozygous, most heterozygous variant sites were converted to homozygous sites using the log-likelihood ratios of reference to alternative genotype calls (Cook et al., 2016). When the log-likelihood ratio was < −2 or > 2, heterozygous genotypes were converted to reference genotypes or alternative genotypes, respectively. All other SNVs with likelihood ratios between −2 and 2 were left as heterozygous variants. Further details can be found in the GitHub repository (https://github.com/AndersenLab/wi-gatk). After variant calling, the following filters were applied with GATK to keep only high-quality variants: read depth (FORMAT/DP > 5); variant quality (INFO/QUAL > 30 and quality by depth INFO/QD > 20); and strand bias (INFO/FS < 100 and INFO/SOR < 5). Variant sites that have a missing genotype in more than 95% of samples or are heterozygous in more than 10% of samples were also removed. With the high-quality set of variants, we ran the concordance-nf pipeline to compare *C. elegans* strains isolated in this study and previously described strains (Cook, Zdraljevic, Roberts, & Andersen, 2017; Cook et al., 2016; Hahnel et al., 2018; Lee et al., 2021). We classified two or more strains as the same isotype if they shared >99.97% SNVs. If a strain did not meet this criterion, we classified it a unique isotype. All genetic analyses in this paper were done on the isotype level. We refer to the final isotype-level VCF as the “isotype VCF”.

### Phylogenetic analyses

We characterized the relatedness of the *C. elegans* isotypes using QuickTree (v2.5) software (Howe, Bateman, & Durbin, 2002). To construct the unrooted tree that includes 540 isotypes (Supplemental Figure 6), we used SNVs from the “isotype VCF” that were converted to the PHYLIPformat (Felsenstein, 1993) using the vcf2phylip.py script (Ortiz, 2019). This tree was visualized using the ggtree (v1.10.5) R package (Yu, Smith, Zhu, Guan, & Lam, 2017). To construct the unrooted trees of 163 Hawaiian isotypes (Supplemental Figure 5), we further pruned the “isotype VCF” by filtering to biallelic SNVs only and removing sites in linkage disequilibrium (LD) using the PLINK (v1.9) commands (--snps-only --biallelic-only --indep-pairwise 50 10 0.8). This --indep-pairwise command uses 50 marker windows and greedily prunes variants from this window that have r^2^ values greater than the threshold (0.8) until no such pairs remain. Then, the sliding window steps forward 10 markers and repeats the process. We also used various pairwise r^2^ thresholds (0.8, 0.6, 0.2, and 0.1) to explore the effects of LD pruning on tree topology (Supplemental Figure 12).

### Population genetic statistics

Genome-wide nucleotide diversity (pi) and Tajima’s D were calculated using the vcftools (v0.1.15) software. We calculated these statistics separately for the 163 Hawaiin isotypes and the 377 non-Hawaiian isotypes by subsetting the full “isotype VCF”. We then calculated genome-wide pi along sliding windows with a 10 kb window size and a 1 kb step size using the (--window-pi 10000 --window-pi-step 1000) commands for both groups. We also calculated Tajima’s D with a 10 kb window size without sliding using the (--TajimaD 10000) command for both groups.

### Principal components analysis and population structure

The smartpca executable from EIGENSOFT (v6.1.4) was used to perform principal components analysis (PCA) (Price et al., 2006). We performed this analysis with the “isotype VCF” that was subset to the 163 Hawaiian isotypes then filtered to biallelic snps only using the PLINK (v.1.9) commands (--snps-only --biallelic-only) and LD pruned with the command (--indep-pairwise 50 10 0.1). We ran smartpca with and without removing outlier isotypes to analyze the population structure among the Hawaiian isotypes. When analyzing the population without removing outlier isotypes, we used the following parameters: altnormstyle: NO, numoutevec: 50, familynames: NO, numoutlieriter: 0. When analyzing the population with outlier isotype removal, we set numoutlieriter to 15. We performed hierarchical cluster analysis on the significant eigenvectors in R using the stats package *hclust* function with the “average” agglomeration method and cut the tree with the cutree function and “k = 7” (R Core Team, 2020). Isotypes were assigned to genetic groups based on the clusters.

### Genotype-environment association and local adaptation

We used two GEA methods (BayPass and GWA) to scan the genome for signatures of local adaptation. Prior to performing GEA with BayPass and GWA, we further pruned the “isotype VCF” by filtering to biallelic SNVs only, removing sites in LD, and setting a minor allele frequency cutoff using the PLINK (v1.9) commands (--snps-only --biallelic-only --indep-pairwise 50 1 0.8 --maf 0.1). We used BayPass (v2.2) (Gautier, 2015) to perform GEA with SNV data from isotypes that were found in 3 km sampling clusters that contained at least three isotypes. This filtering strategy resulted in 142 isotypes from 13 sampling clusters for use in BayPass. To generate a scaled population covariate matrix (omega matrix), we first ran BayPass with the core model using 13 populations (3 km sampling clusters) and a subsampled dataset of 5000 biallelic SNVs from outside annotated gene coding regions generated using the PLINK (v1.9) commands --exclude and --thin-count 5000 The omega matrix was then used to explicitly account for the covariance structure in population allele frequencies resulting from the demographic history of the populations in subsequent BayPass runs. We reran the BayPass core model using the full set of biallelic SNVs, 13 populations, and the omega matrix. The BayPass core model outputs XtX statistics for each marker that can be used to identify differentiation caused by selection rather than other processes. We considered XtX statistic values as suggestive of local adaptation if they were among the top 0.1% of genome-wide XtX statistic values. We then ran BayPass again using the standard covariate model with biallelic SNV data, 13 populations, the omega matrix, and eight environmental variables as covariates (altitude, mean annual air temperature, mean annual surface temperature, mean annual rainfall, mean annual soil moisture, mean annual leaf area index, latitude, and longitude). We considered Bayes Factors (BFs) above 20 as evidence of a significant genotype-environment association. We also performed GEA with the NemaScan pipeline, a GWA tool specifically designed for *C. elegans*, which is available at https://github.com/AndersenLab/NemaScan (Widmayer, Evans, Zdraljevic, & Andersen, 2021). NemaScan uses both the --mlma-loco and --fastGWA-lmm-exact functions from GCTA software to perform rapid GWA (Yang, Lee, Goddard, & Visscher, 2011). The --mlma-loco function accepts a limited sparse kinship matrix composed of all chromosomes except the chromosome containing the tested marker (LOCO = “leave one chromosome out”) and the --fastGWA-lmm-exact accepts a full sparse kinship matrix specifically calculated for inbred model organisms. These functions can account for population structure in the mapping sample but use a different strategy than BayPass. To determine significant markers, we used eigen decomposition of the kinship matrix to correct for the number of independent tests in each mapping, as was described previously (Zdraljevic et al., 2019).

To identify genomic regions of interest that contained markers significantly associated with environmental parameters, we grouped markers into a single region if they were within 1 kb of each other and were above the significance threshold. We then expanded that region of interest to include the portions of the genome containing 150 markers to the left and right of its original range. Ultimately, we identified overlapping regions of interest for the two GEA methods using the bedtools (v2.30) intersect command.

### Data description

Supplemental File 1: Sampling data for every Hawaiian collection used to generate Figure 1.

**Figure 1.**
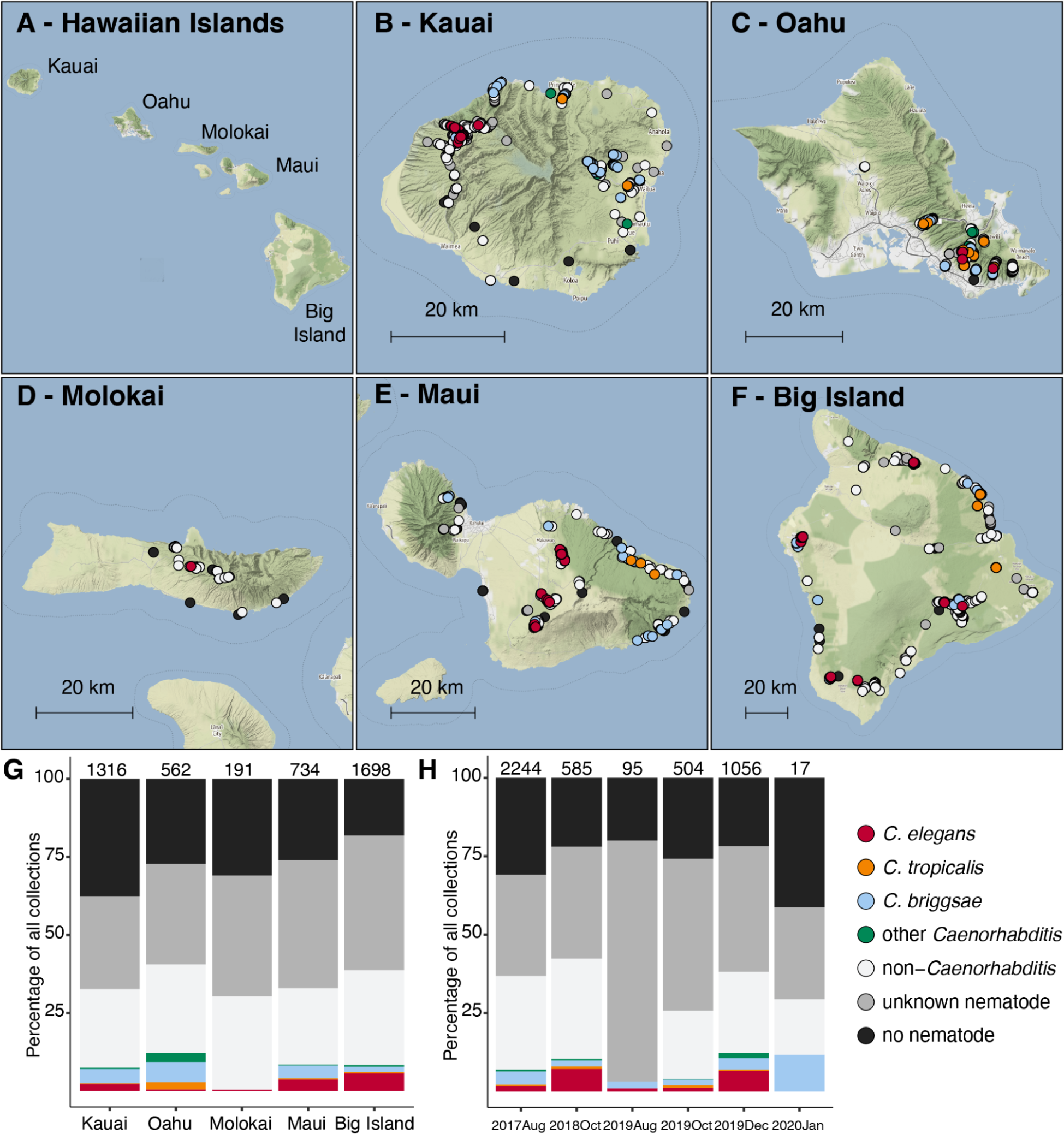
Geographic and temporal distribution of sampling sites across the Hawaiian Islands. (A) An overview of the Hawaiian Islands. (B-F) Detailed views of each of the islands sampled. Circles indicate sampling sites and are colored according to the legend below. We categorized nematodes as “other-*Caenorhabditis*” if they did not belong to one of the three selfing *Caenorhabditis* species and “non-*Caenorhabditis*” if their ITS2 region aligned to genera other than *Caenorhabditis* or if the ITS2 region failed to amplify but the 18S region did. We categorized nematodes as “unknown nematodes” if we could not extract high-quality genomic DNA or amplify either region by PCR (see, Materials and Methods). For sampling sites where multiple collection categories apply (n = 733), the site is colored by the collection category shown in the legend from top to bottom, respectively. (G-H) The percentage of each collection category is shown by island (G) or collection project (H). Bars are colored according to the legend on the right and the total number of samples for each category are shown above the bar.

Supplemental File 2: Collection class frequencies for each land cover type within habitat class used to generate Figure 2.

**Figure 2.**
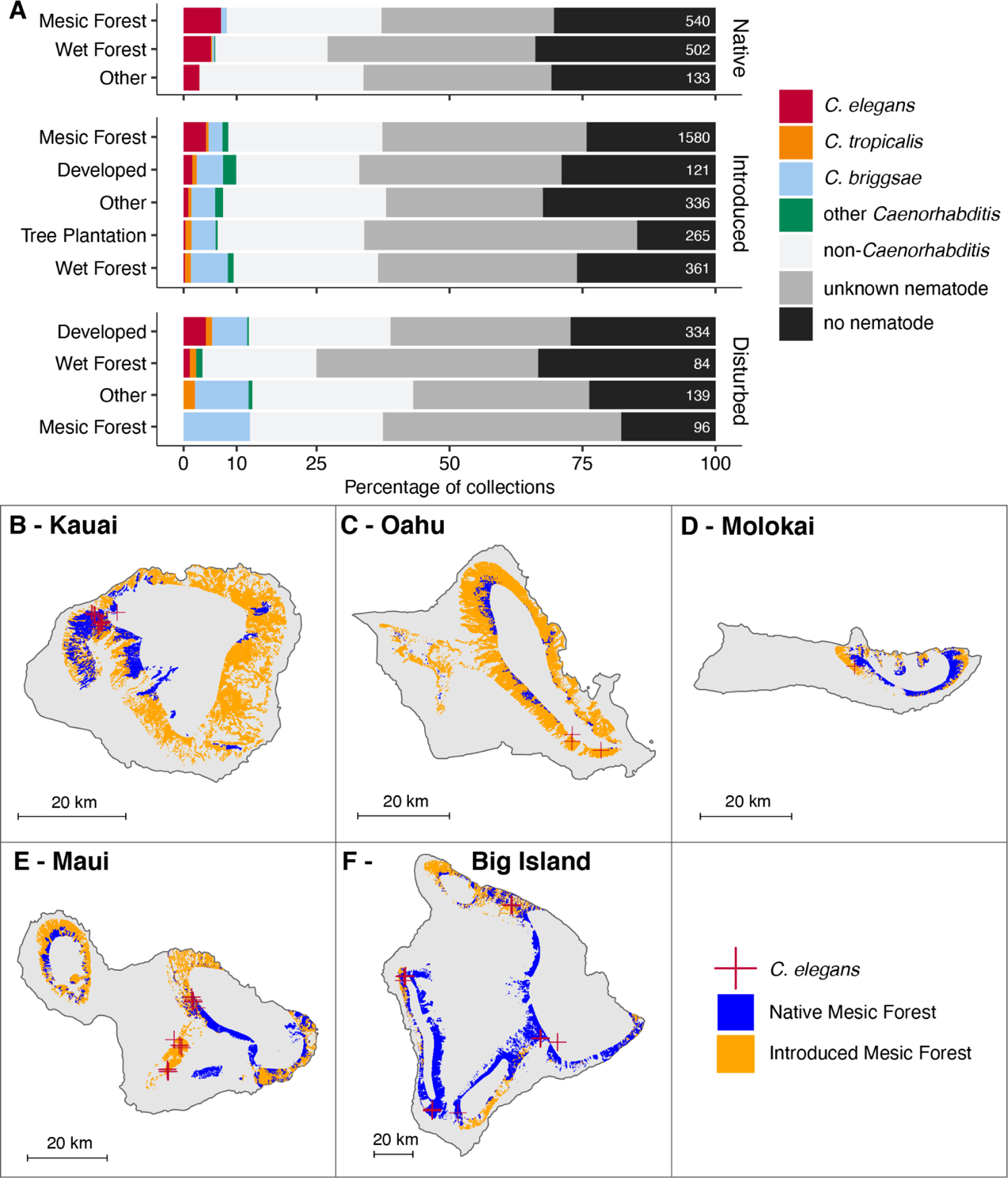
Land cover enrichment among selfing *Caenorhabditis* nematodes. (A) The percentage of each sampling category is shown by land cover type. The land cover types are organized by native, introduced, and disturbed habitats. The sampling categories are colored according to the legend at the right, and the total number of samples for each substrate are shown on the right side of the bars. (B-F) Land cover maps for each of the five Hawaiian Islands sampled in this study. The sample locations where *C. elegans* were found are shown as red crosses, and native and introduced mesic forest land covers are shaded blue and orange respectively.

Supplemental File 3: Collection class frequencies for each substrate type used to generate Figure 3A.

Supplemental File 4: Environmental parameter values for samples used to generate Figure 3B-G, Supplemental Figure 2, and Supplemental Figure 3.

**Figure 3.**
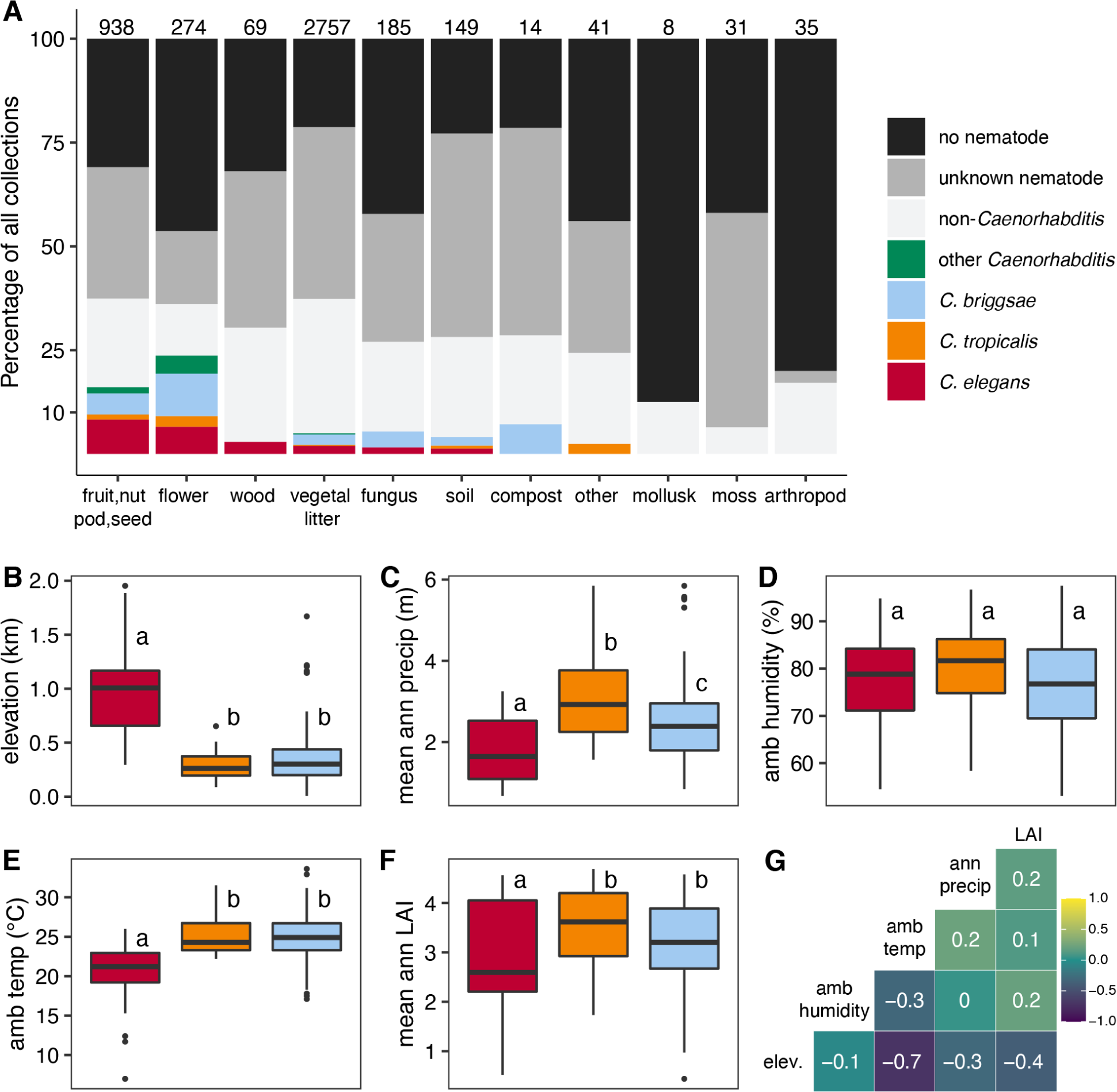
Niche differentiation among selfing *Caenorhabditis* nematodes. (A) The percentage of each sampling category is shown by substrate type. The sampling categories are colored according to the legend on the right, and the total number of samples for each substrate are shown above the bars. (B-F) Environmental parameter values; elevation, mean annual precipitation, *in situ* ambient humidity, *in situ* ambient temperature, and mean annual leaf area index (LAI) for sites where *Caenorhabditis* species were isolated. Tukey box plots are plotted by species (red is *C. elegans*, orange is *C. tropicalis*, blue is *C. briggsae*) for each environmental parameter; points above or below whiskers indicate outliers. Letters above the boxes summarize the statistical significance of comparisons between the species shown. Species with a different letter are significantly different; species with the same letter are not significantly different. Comparisons were made using a Kruskal-Wallis test and Dunn’s test for multiple comparisons with *p*-values adjusted using the Bonferroni method. (G) A correlation matrix for the continuous environmental parameters shown. The parameter labels for the matrix are printed on the diagonal, and the Pearson correlation coefficients are printed in the cells. The color scale also indicates the strength and sign of the correlations shown in the matrix.

Supplemental File 5: Genetic group assignments and PC values for all non-outlier isotypes used to make Figure 4A-B, Supplemental Figure 11, and Supplemental Figure 12D.

**Figure 4.**
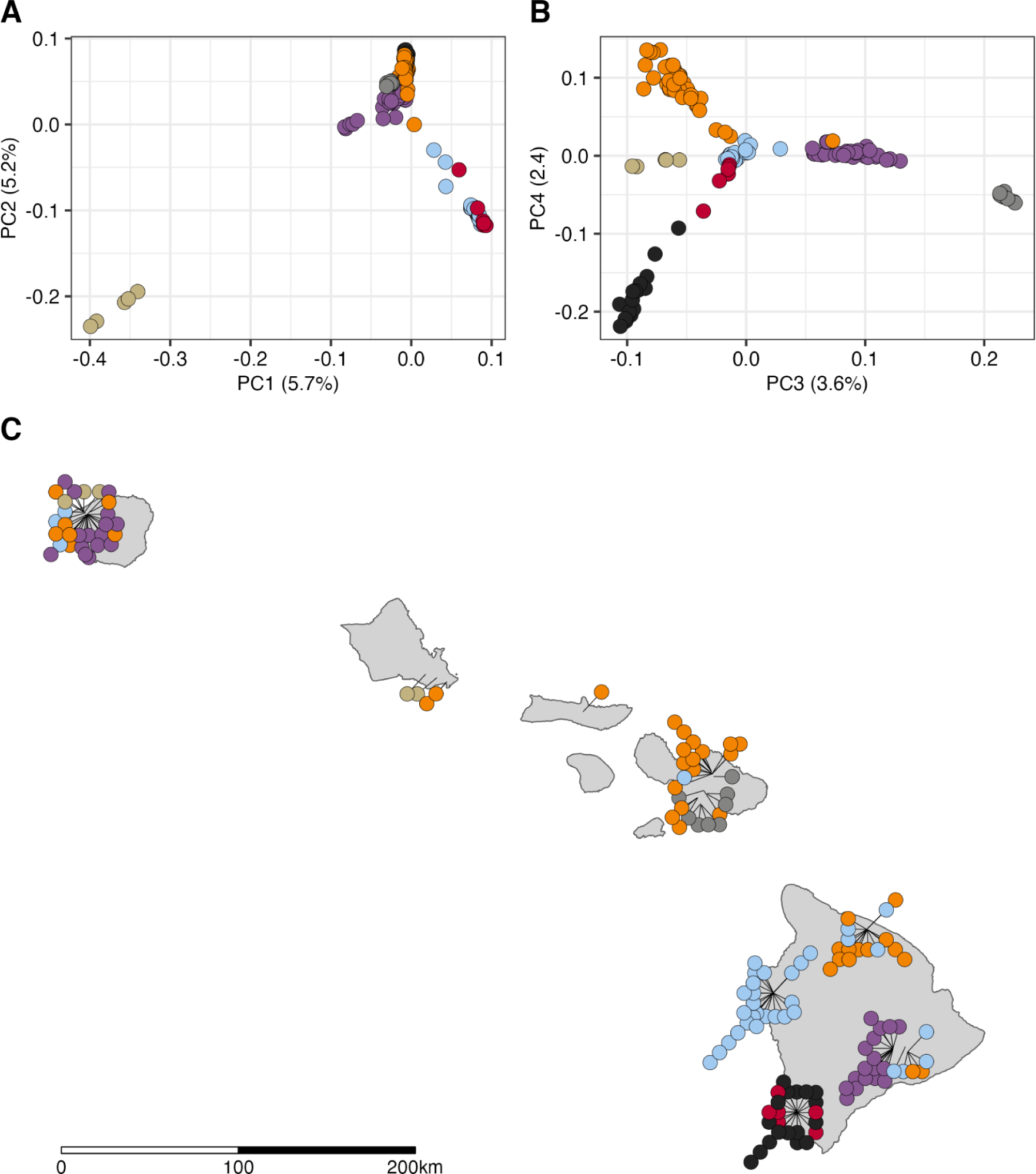
*C. elegans* genetic structure in Hawaii. (A-B) Plots show major axes of variation derived from principal components analysis (PCA) of the genotype covariance matrix of 163 Hawaiian isotypes. (A) The first two axes of variation are plotted (PC1 and PC2). (B) The third and forth axes of variation are plotted (PC3 and PC4). (A-B) The points indicate individual isotypes and are colored by genetic group assignments obtained from hierarchical clustering of eigenvalues. Only 149 of 163 isotypes are shown, 14 outlier isotypes were removed from the PCA (see Materials and Methods). (C) The sampling locations for 144 of 163 Hawaiian isotypes are plotted on Hawaii. Each circle represents a single isotype and is colored by genetic group assignment. The 14 isotypes that are PCA outliers and five isotypes without location data are not shown.

Supplemental File 6: Geographic locations of sampling sites for isotypes with genetic group assignments used to make Figure 4C.

Supplemental File 7: BayPass Bayes Factors for SNVs used to generate Figure 5.

**Figure 5.**
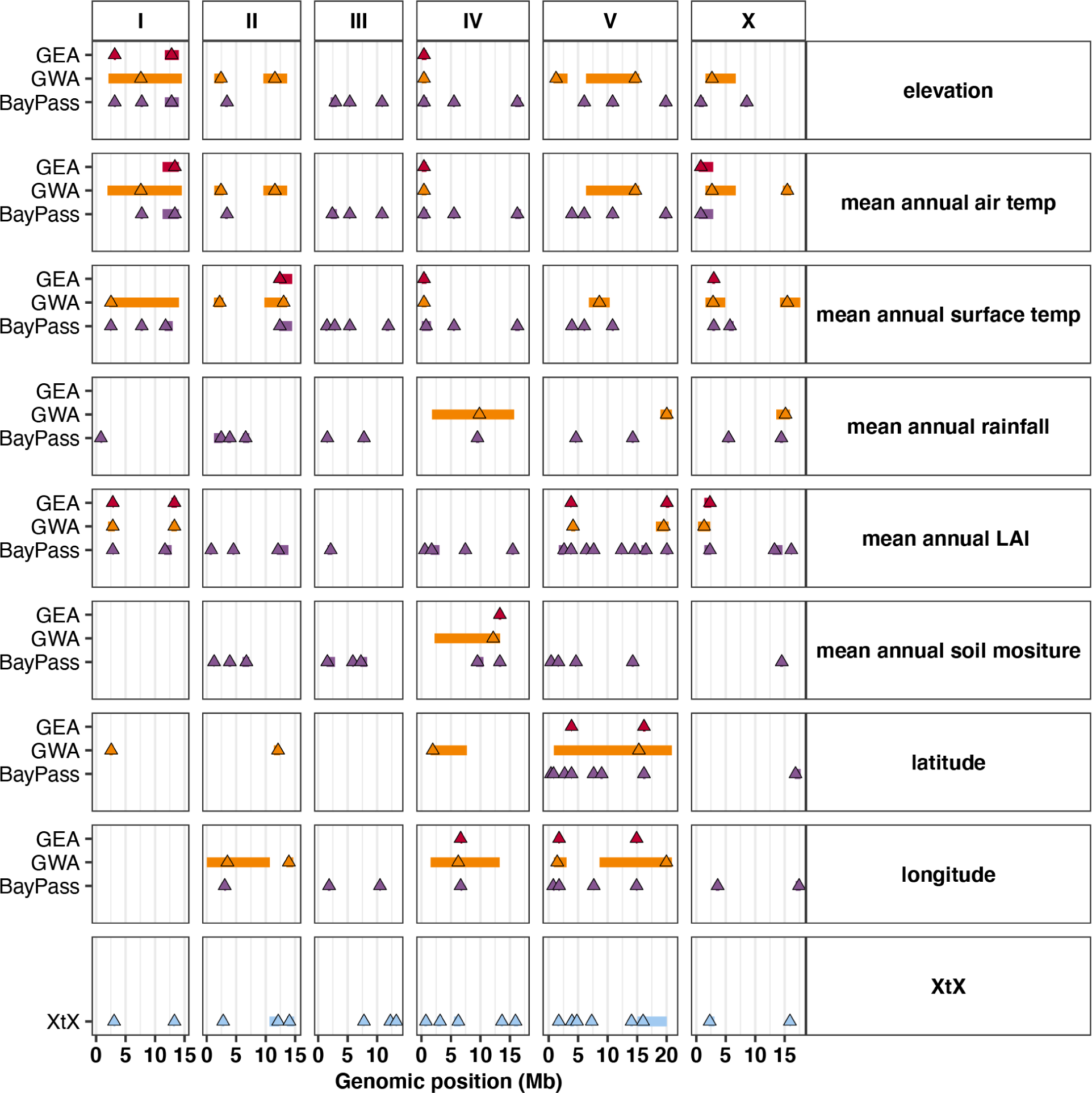
Genetic architecture of local adaptation. Genotype-environment association (GEA) results for BayPass and GWA methods are plotted for eight environmental variables: elevation, mean annual air temperature, mean annual surface temperature, mean annual rainfall, mean annual leaf area index (LAI), mean annual soil moisture, latitude, and longitude. The BayPass XtX statistic is plotted on the bottom facet. Each triangle represents the marker with peak significance value within each region for each method. Regions are shown as rectangles plotted behind the peak marker position and their width is determined as described in the Materials and Methods. The peak markers and regions are colored by their type (red = GEA region, orange = GWA region, purple = BayPass region, blue = XtX region). The GEA regions represent cases where BayPass, GWA, and XtX regions overlap. The overlap for BayPass and GWA are considered within the same environmental variable. If overlap exists, then the smaller of the two overlapping regions is compared to the XtX statistic. If the smaller of the two regions overlaps with the XtX statistic, then the region is determined to be a GEA region and plotted in red for each environmental variable. Genomic position is plotted along the x-axis in megabases and by chromosome.

Supplemental File 8: BayPass XtX statistics for SNVs used to generate Figure 5.

Supplemental File 9: GWA regions used to generate Figure 5.

Supplemental File 10: GEA regions used to generate Figure 5.

Supplemental File 11: Counts for collection classes by project used to generate Supplemental Table 1.

Supplemental File 12: Cohabitation counts for distinct taxa used to generate Supplemental Figure 1.

Supplemental File 13: Sampling frequencies for habitat classes by substrate type used to generate Supplemental Figure 4.

Supplemental File 14: Sampling frequencies for land cover class by substrate type grouped by habitat class used to generate Supplemental Figure 5.

Supplemental File 15: Pairwise geographic distances between distinct collections within each isotype used to generate Supplemental Figure 6.

Supplemental File 16: Geographic coordinates of 3 km diameter sampling clusters and the number of isotypes sampled within each cluster used to generate Supplemental Figure 7.

Supplemental File 17: Sampling frequencies of temporally persistent isotypes over time for each sampling location used to generate Supplemental Figure 8.

Supplemental File 18: Sweep status and number of swept chromosomes for 540 isotypes sampled from around the world used to generate Supplemental Figure 9.

Supplemental File 19: Genome-wide pi calculated for 163 Hawaiian and 377 non-Hawaiian isotypes in sliding 10 kb windows that was used to generate Supplemental Figure 10A.

Supplemental File 20: Genome-wide Tajima’s D calculated for 163 Hawaiian and 377 non-Hawaiian isotypes in 10 kb windows that was used to generate Supplemental Figure 10B.

Supplemental File 21: PC values for all 163 Hawaiian isotypes generated by PCA without outlier removal that was used to generate Supplemental Figure 11.

Supplemental File 22: Genetic group assignments derived from PCA and clustering using different LD thresholds prior to PCA (r^2^ = 0.8, 0.6, 0.2, 0.1) used to make Supplemental Figure 12.

Supplemental File 23: Correlation matrix for significant genetic PCs by continuous environmental parameters.

Supplemental File 24: Continuous environmental parameter values for distinct collections within each genetic group used to generate Supplemental Figure 14.

## Results

### Hawaiian nematode diversity

We collected Hawaiian nematodes on six occasions between August 2017 and January 2020. On each collection trip, we performed multiple sampling trials in different habitats and preferentially sampled substrates known to harbor *Caenorhabditis* nematodes, including rotting fruits, seeds, nuts, flowers, stems, mixed vegetal litter, compost, wood, soil, fungus, live arthropods, and molluscs (Crombie et al., 2019; Ferrari et al., 2017; Schulenburg & Félix, 2017). At each sampling site, we measured various environmental parameters, including ambient humidity and temperature, elevation, and substrate temperature using hand-held devices (Di Bernardo et al., 2021). We sampled from some hiking trails on multiple collection trips, but other trails were only sampled once. Overall, we sampled 4,506 substrates across five Hawaiian Islands (Figure 1) and isolated 7,107 nematodes from 2,400 of these substrates (53.3% success rate for nematode isolation). We attempted to identify these isolates by analysis of the 18S rDNA gene and the Internal Transcribed Spacer (ITS2) region between the 5.8S and 28S rDNA genes (see Materials and Methods) (Barrière & Félix, 2014; K. C. Kiontke et al., 2011). We identified *Caenorhabditis* isolates to the species-level and isolates from other genera to the genus-level. In total, we identified 23 distinct taxa across the Hawaiian Islands, including five *Caenorhabditis* species that were found at different frequencies among the 4,506 substrates sampled: *C. briggsae* (3.6%), *C. elegans* (3.6%), *C. oiwi* (0.68%), *C. tropicalis* (0.58%), and *C. kamaaina* 0.09%) (Supplemental Table 1). We also collected 13 samples (0.29%) harboring 31 isolates with ITS2 regions that were most closely related to either *C. plicata* or *C. parvicauda* but at such low identity that we suspect the isolates belong to one or more new *Caenorhabditis* species. However, all of these isolates perished before they could be cryopreserved so we have classified them as *“Caenorhabditis* spp.”.

Our collection data suggest that nematode diversity is similar across the five sampled islands. The number of taxa identified on each island scaled with the number of samples collected and ranged from 20 taxa on the Big Island to just three taxa on Molokai. However, we did not detect an enrichment of diversity on any particular island when considering the number of taxa relative to the number of genotyped isolates from the island (Fisher’s Exact Test; *p* > 0.05 for all island comparisons). We also looked for differences in diversity at a finer geographic scale (∼3 km) by comparing diversity among various sampled trails. We isolated between zero and 11 taxa per trail, but we found no evidence of enrichment on any particular trail (Fisher’s Exact Test; *p* > 0.05 for all trail comparisons). In rare cases, we observed nematode diversity at the substrate scale (∼10 cm). Among the 4,506 substrates sampled, we observed 46 instances of distinct taxa cohabitating on the same substrate, a finding consistent with previous surveys of nematodes (Crombie et al., 2019; Félix et al., 2013; Petersen et al., 2014; Richaud et al., 2018). Among these samples, 21 contained more than one *Caenorhabditis* species (Supplemental Figure 1). These patterns of *Caenorhabditis* nematode diversity suggest that species within the genus could compete for resources across multiple spatial scales. Given the high frequencies of colocalization among *Caenorhabditis* species, we searched for observable differences in environmental niche preferences among them.

### Caenorhabditis niche specificity

To characterize the niche preferences of *Caenorhabditis* nematodes, we used publicly available geospatial data to identify habitats where they are found most frequently. We used a GIS map of habitat status to assign each sampling site one of three habitat conditions: native, introduced, or disturbed (see Materials and Methods) (Jacobi et al., 2017). Native habitats are dominated by native Hawaiian plant communities; introduced habitats contain a mixture of native and introduced species; and disturbed habitats are impacted by agriculture or urban development. Overall, we found *Caenorhabditis* nematodes were enriched in disturbed habitats (74 of 653 11.3%) relative to native habitats (78 of 1175 6.6%) but not relative to introduced habitats (221 of 2663 8.3%) (Fisher’s Exact Test; disturbed vs native *p* = 0.0044, disturbed vs introduced *p* = 0.10). The pattern of habitat enrichment was strikingly different for *C. elegans* relative to the other selfing species. Whereas *C. briggsae* and *C. tropicalis* were enriched in disturbed habitats relative to native habitats, *C. elegans* was enriched in native habitats (68 of 1175 5.8%) relative to introduced (74 of 2263 2.8%) or disturbed habitats (15 of 653 2.3%) (Fisher’s Exact Tests, *p* < 0.0025) (Figure 2A). We also used land cover GIS data to determine whether *Caenorhabditis* nematodes were associated with a particular land cover class (Jacobi et al., 2017). We found no evidence of *Caenorhabditis* enrichment for any particular land cover within native, introduced, or disturbed habitat classes. However, within native and introduced habitats *C. elegans* were found most frequently in mesic forests compared to the other land covers. Hawaiian mesic forests have moderate amounts of rainfall (1200 to 2500 mm annually) and are found on leeward and windward sides of the islands in lowland or in montane-subalpine zones (Cuddihy, Pratt, & Stone, 1990). Among introduced habitats, *C. elegans* were significantly enriched in mesic forest relative to all other land covers except developed (Fisher’s Exact Test; *p* < 0.012) (Figure 2A). Notably, the number of samples we collected from introduced habitats with developed land cover was small (n = 121) and both samples that contained *C. elegans* (n = 2) were taken from the roadside adjacent to mesic forest land cover. Taken together, these data suggest that *C. elegans* niche preferences might be different from other selfing *Caenorhabditis* species in Hawaii. *C. elegans* seems to prefer native mesic forest habitats, but *C. briggsae* and *C. tropicalis* are more frequently found in disturbed and introduced habitats and at similar frequencies across land covers within these habitats (Figure 2A).

On the Hawaiian Islands, elevation and tradewinds influence species assemblages and land covers by causing gradients in various environmental parameters, including temperature, and rainfall (Lowe, Ball, Reeves, Amidon, & Miller, 2020). To further characterize Hawaiian *Caenorhabditis* niche preferences, we measured various environmental parameters at each sampling site *in situ*, including elevation, substrate temperature, ambient temperature, and ambient humidity. We also obtained measurements of additional continuous environmental parameters from geospatial models by extracting data associated with the location of each sampling site (see Materials and Methods, Supplemental Figure 2,3) (Frazier et al., 2016; Thomas W. Giambelluca et al., 2013; T. W. Giambelluca et al., 2014). To reduce the dimensionality of these data, we choose to focus on just five of the nine variables by removing the variable with the most missing data and the correlation was greater than 0.7 (Figure 3B-G). We observed that on average, *C. elegans* were found at cooler, higher elevations in less densely vegetated and drier regions than the other selfing *Caenorhabditis* species (Kruskal-Wallis and Dunn’s post-hoc test *p* < 0.05; ambient temperature, elevation, mean annual leaf area index, ambient humidity, mean annual precipitation). We sampled *C. elegans* from surprisingly cold substrates; three high-elevation *C. elegans*-positive collections from Maui (elevation > 1400 m) made in December 2019 were temperature outliers among all selfing species. One of these substrates was collected at an ambient temperature of 7.0°C, with a substrate temperature of 3.9°C. We did not observe differences between *C. briggsae* and *C. tropicalis* species for the majority of environmental parameters, although all three species were differentiated from each other with respect to mean annual precipitation (Figure 3C). On average, *C. elegans* was found at sites with the lowest precipitation values relative to *C. briggsae* and *C. tropicalis* (Kruskal-Wallis and Dunn’s post-hoc test *p* < 0.05 for all comparisons). These trends suggest that wild *C. elegans* tend to prefer high altitude, cooler, native dominated, mesic forest habitats. *C. briggsae* and *C. tropicalis* are typically found at lower elevations in warmer, wetter, less native habitats. Consistent with these patterns, the cohabitation frequency between *C. tropicalis* and *C. briggsae* (2.6%, 5 of 190 collections) is higher than the cohabitation frequency between *C. elegans* and *C. briggsae* (0.9%, 3 of 326 collections) or *C. elegans* and *C. tropicalis* (0%, 0 of 188 collections). Moreover, the three samples harboring both *C. elegans* and *C. briggsae* were collected between 650 and 800 meters, near the lower range of *C. elegans* elevations and the upper range of *C. briggsae* elevations.

We also classified substrate types at each sampling location to explore possible preferences among the three selfing *Caenorhabditis* species. We isolated *Caenorhabditis* nematodes at higher frequencies from flower (65 of 274 or 23.7%) and fruit substrates (151 of 938 or 16.0%) than any other category except compost (Fisher’s Exact Test, *p* < 0.05) (Figure 3A). Notably, the sample size for wood, compost, and other substrates was low, which limits our power to detect *Caenorhabditis* preferences for these substrates. These trends underscored known preferences for decomposing flower and fruit substrates identified by previous surveys of wild *Caenorhabditis* nematodes in tropical regions (Crombie et al., 2019; Félix et al., 2013; Ferrari et al., 2017). Although *Caenorhabditis* nematodes are associated with invertebrates in the wild (K. Kiontke & Sudhaus, 2006), we did not isolate them from either mollusks or arthropods, but our sample size on these substrates was also small (Figure 3A). We did not observe differences in the patterns of substrate enrichment among the selfing species; all species were found more frequently on fruit and flower substrates than vegetal litter (Fisher’s Exact Test, *p* < 0.05) (Figure 3A). This observation differs from our initial survey of Hawaiian nematodes, where we did not identify a significant substrate enrichment for *C. elegans* (Crombie et al., 2019). Importantly, the *C. elegans* enrichment in native habitats described above is not caused by oversampling of fruit or flower substrates in native habitats. In fact, we sampled fruit and flower substrates less frequently in native habitats than introduced or disturbed habitats (Supplemental Figure 4). However, it is possible that unequal sampling of preferred substrates among different land covers could cause the enrichment of *C. elegans* in mesic forests (Supplemental Figure 5). Additional sampling of diverse substrate types across different land covers will be needed to address this possibility.

Overall, the sampling data reveal that *C. elegans* niche preferences are likely different from *C. briggsae* and *C. tropicalis* but also reveal considerable variation within the niche for each species. For example, although *C. elegans* are typically found in native habitats, a number of *C. elegans* were isolated from disturbed habitats in developed areas. Moreover, although *C. elegans* was on average found at cooler, higher elevations than other *Caenorhabditis* species, we isolated the species across a wide range of temperatures (7 - 26°C) and elevations (295 - 1950 meters). We were curious whether the genetic diversity of Hawaiian *C. elegans* might be associated with variation in niche parameters on the islands. To explore this possibility, we sequenced the genomes of the *C. elegans* strains we had isolated.

### Genetic diversity of Hawaiian C. elegans

We sequenced the genomes of the 464 extant Hawaiian *C. elegans* strains to high coverage (median 27x) and identified single nucleotide variants (SNVs) and small indel variants in each genome relative to the N2 reference genome (36 strains were lost before cryopreservation). In *C. elegans*, some wild strain genome sequences are often nearly identical because of the high rates of self-fertilization in the species. To reduce the number of invariant genomes in our analyses, we measured the concordance among all wild strain pairs and grouped strains sharing >99.97% of SNVs into a single genome-wide haplotype we refer to as an “isotype” (see Materials and Methods) (Andersen et al., 2012). Using this strategy, we identified 143 Hawaiian isotypes among the 464 wild strains we sequenced. We expanded the sample to 163 Hawaiian isotypes with 20 additional isotypes that were either sampled prior to 2017 or more recently by our collaborators. The vast majority of the Hawaiian isotypes comprise strains that were sampled from the same substrate (134 of 163, 82%). Among the 29 isotypes sampled from multiple substrates, the median sampling distance between substrates that have the same isotype was 35.9 meters (Supplemental Figure 6). Notably, only three isotypes comprised strains sampled more than 500 meters from one another over multiple years. For example, isotype XZ1513 was first sampled in 2014 from the island of Kauai and then sampled again in 2018 from a location more than six kilometers away from the original sampling location.

We applied hierarchical clustering methods to assign *C. elegans* positive samples to one of 21 discrete three kilometer diameter sampling locations across the islands (see Materials and Methods). The vast majority of Hawaiian isotypes comprised strains that were sampled in close proximity to one another. Specifically, 157 of the 163 isotypes were never sampled from more than one of the 21 sampling locations, two isotypes were found in two sampling locations, and four isotypes could not be clustered because GPS positions were not available (Supplemental Figure 7). Similar to another longitudinal survey of genetic diversity in *C. elegans (Richaud et al., 2018)*, we identified the same isotypes that persisted at sampling locations over multiple years (Supplemental Figure 8). However, unlike previous studies, we never observed a single isotype that persisted at high frequency within a sampling location. In each of the five sampling locations where isotypes persisted, the average persistence time between the first and last isolation was 684 days, and in all cases, the relative abundance of the isotype in the sampling location varied substantially over time. We did not observe obvious seasonal variation in *C. elegans* abundance or relative abundance of isotypes at sampling locations, but these patterns might be observable with additional sampling at regular intervals. In general, we found a high degree of genetic diversity within sampling locations relative to longitudinal surveys of diversity in Europe. Among the 10 sampling locations where we collected at least five samples, we found an average of 0.93 isotypes per sample. By comparison, the number of isotypes sampled from non-Hawaiian sampling locations is often much lower with 0.29 isotypes per sample in France and 0.27 isotypes per sample in Germany (Andersen et al., 2012; Cook et al., 2016; Lee et al., 2021; Petersen et al., 2014; Richaud et al., 2018). The identification of more isotypes per sample in Hawaiian is not because we genotyped more individuals from a given sample. In fact, the number of individuals genotyped per sample was on average twice as high in the European locations (∼5 individuals per sample) as it was in our Hawaiian locations (∼2.5 individuals per sample). Taken together, these data suggest that genetic diversity at the local scale (1 to 10 km^2^) is higher on the Hawaiian Islands than it is in France and Germany.

Across a total of 163 Hawaiian isotypes, we identified 2.6 million SNVs and 507,680 small indels, which is twice the number of variants found among the 377 non-Hawaiian isotypes known at the time of this study (Cook et al., 2017) (release 20210121). Moreover, we found that approximately 60% of all SNVs and indels described in *C. elegans* are only found in Hawaiian isotypes and that nucleotide diversity (pi) is almost three-fold higher in the set of Hawaiian isotypes than it is among non-Hawaiian isotypes (Hawaii pi = 0.0031; non-Hawaiian pi = 0.0012; Supplemental Figure 9). Consistent with previous analyses of genetic diversity in *C. elegans*, we detected higher levels of diversity in both Hawaiian and non-Hawaiian samples on chromosome arms relative to centers, a pattern that is likely caused by higher rates of recombination and therefore reduced levels of background selection on chromosome arms (Andersen et al., 2012; Crombie et al., 2019; Cutter & Payseur, 2003; Rockman, Skrovanek, & Kruglyak, 2010). We also detected multiple localized spikes in pi with values exceeding six times the genome-wide average for both Hawaiian and non-Hawaiian samples. These spikes in diversity often overlap with spikes in Tajima’s D and likely reflect hyperdivergent regions that are theorized to be maintained in *C. elegans* by long-term balancing selection (Lee et al., 2021; Thompson et al., 2015). We suspect that the remarkable genetic diversity in the Hawaiian isotypes relative to the non-Hawaiian isotypes is at least partially caused by the near absence of selective sweeps in the Hawaiian isotypes. These megabase-scale selective sweeps are thought to have purged genetic diversity from many non-Hawaiian isotypes across the centers of chromosomes I, IV, V, and the left arm of chromosome X (Andersen et al., 2012). Indeed, we found that 93% (351 of 377) of non-Hawaiian isotypes have at least one swept chromosome compared to just 4% (6 of 163) of Hawaiian isotypes, and this pattern likely contributes to the high degree of differentiation for most Hawaiian isotypes (Supplemental Figure 10). Consistent with the theory that sweeps offer fitness advantages in human-associated habitats, four of the six isotypes with selective sweeps were isolated from disturbed or introduced habitats with developed land covers. Moreover, two of those isotypes (ECA923 and ECA928) were isolated from a backyard garden in downtown Honolulu. However, one isotype (XZ1515) with a global swept haplotype was isolated from a native mesic forest habitat in Kauai so globally swept haplotypes are also present in at least some native habitats in Hawaii.

### Genetic structure of Hawaiian C. elegans

We examined the genetic structure among the 163 Hawaiian isotypes using principal components analysis (PCA) (Price et al., 2006). We found six significant principal components (PCs) that together explain 21% of the genetic variation within the Hawaiian isotypes. We then applied hierarchical clustering to these axes of variation and subdivided the Hawaiian sample into seven genetically distinct groups (Figure 4, Supplemental Figures 11, 12). To explore whether these genetic groups were associated with geography, we plotted isotype sampling coordinates onto a map of Hawaii and colored them by their genetic group assignments (Figure 4). We found that some genetic groups were widely distributed across the islands, and others were restricted to individual sampling locations. For example, the orange group consisted of 42 isotypes that were sampled from all five islands. By contrast, the red and black groups consisted of six and 17 isotypes, respectively, and were each sampled exclusively from Manuka State Wayside Park located on the south side of the Big Island. We next tested whether the PCs were correlated with either spatial coordinates or continuous environmental parameters. We found that the PC1, which explained 5.7% of the total genetic variation, was significantly correlated with both longitude and latitude, suggesting that genetic variation is at least partially associated with geography (Supplemental Figure 13). We also detected significant correlations between other axes of genetic variation and continuous environmental parameters. For example, PC3 is positively correlated with altitude and negatively correlated with various measures of temperature at isotype sampling sites. Moreover, PC4 is positively associated with measures of moisture at isotype sampling sites. Together, these data suggest that genetic variation among the Hawaiian isotypes could be partially explained by local adaptation to environmental conditions at sampling locations, although these correlations might also be caused by neutral processes. The purple group offers another compelling indication that local adaptation contributes to the genetic variation we observe on the Hawaiian Islands. It comprises 34 isotypes that were sampled from two locations separated by over 500 km, the first on Kauai (15 isotypes) and the other on the Big Island (17 isotypes) (Figure 4). Despite the extreme distance separating these sampling locations, they have similar environments with respect to elevation, temperature, and moisture (Supplemental Figure 14). Moreover, both locations contain high-elevation, native ‘Oh’a and Koa mesic forest habitats, further suggesting that individuals within this genetic group might have been selected in this type of environment.

### Signatures of local adaptation

Local adaptation can occur in response to different selection pressures caused by the environment. For instance, allele frequencies will correlate with selective features of the environment when the selection pressure can counteract homogenizing forces like migration (Haldane, 1948; Lenormand, 2002; Nagylaki, 1975). GEA methods have the potential to detect these signatures of local adaptation and help elucidate their genetic basis in natural populations. We applied two separate GEA methodologies (BayPass and genome-wide association, GWA) to detect signatures of local adaptation to various environmental parameters, including elevation, along with annual mean measures of ambient temperature, substrate temperature, rainfall, soil moisture, and leaf area index (see Materials and Methods). We also explored associations with latitude and longitude because selective pressures associated with geographic location could also underlie signatures of local adaptation. Using the NemaScan GWA pipeline, we found 39 regions of the genome that were associated with environmental or geographic parameters (Figure 5). The BayPass genome-wide scan revealed 108 regions of the genome that were associated with environmental or geographic parameters (Figure 5). On average, the BayPass regions were smaller than the regions identified by GWA. We explored the consensus between these two GEA methods with respect to mappings for each environmental or geographic parameter and found a total of 35 regions that overlapped between the two methods. We refer to these 35 regions as method overlap regions. We also used the XtX statistics calculated by BayPass to scan for genomic regions that appeared to be adaptively differentiated when controlling for the covariance structure in population allele frequencies resulting from the demographic history of the populations (see Materials and Methods) (Gautier, 2015; Günther & Coop, 2013). This scan identified 21 regions of the genome consistent with adaptation to local environments. We then determined that 20 of the 35 method overlap regions fall into 11 XtX regions, we refer to these 20 regions as GEA regions (Figure 5). We prioritized these 20 GEA regions to explore the genetic basis of local adaptations to environmental parameters within the Hawaiian isotypes. In general, we observed that the genetic architectures of environmental associations were similar for correlated variables. For example, an identical region on chromosome IV at 0.8 megabases is associated with elevation and mean annual air and surface temperatures that have pairwise Pearson’s correlation coefficients greater than 0.95 or less than -0.97 (Supplemental Figure 2). Of the 20 GEA regions, 18 are distinct, because of the overlap on chromosome IV described above. Interestingly, all but one of the 18 distinct GEA regions fall into hyperdivergent regions (Lee et al., 2021; Thompson et al., 2015). The hyperdivergent regions are maintained by balancing selection and are enriched for genes related to environmental sensing, xenobiotic detoxification, and pathogen resistance.

Therefore, the signatures of local adaptation we observe in the Hawaiian strains match the expectation that hyperdivergent regions are important for the adaptation of *C. elegans* to its local environment.

## Discussion

This study represents the most detailed survey of *C. elegans* genetic diversity on the Hawaiian Islands to date. Similar to previous studies, we found that Hawaiian strains harbor high levels of genetic diversity relative to strains found in most other sampling locations, particularly locations in Europe (Andersen et al., 2012; Lee et al., 2021; Richaud et al., 2018; Rockman & Kruglyak, 2009). The *C. elegans* niche on the Hawaiian Islands is higher in elevation, cooler, drier, and less impacted by introduced plant communities and human disturbance than other selfing *Caenorhabditis* species. Importantly, our longitudinal sampling strategy uncovered multiple sites across various islands where *C. elegans* can be sampled reliably over time, enabling more detailed examinations of the temporal patterns of genetic variation in the species with additional sampling in the future. Furthermore, our quantitative measures of the *C. elegans* niche will allow for targeted exploration of new sampling locations on Hawaiian Islands and perhaps other Pacific Islands. In general, surveys of other islands in the Pacific and the greater Pacific Rim will help locate the geographic origin of the species and further our understanding of the evolutionary forces shaping the current patterns of genetic diversity.

### Forces contributing to the genetic differentiation in Hawaii

We detected seven genetically distinct groups within our current sample of non-redundant Hawaiian strains. What factors account for the substantial genetic differentiation that we observed across the Hawaiian Islands? In other selfing species such as *Arabidopsis thaliana*, regional patterns of differentiation have been attributed to isolation by distance (IBD) (Manzano-Piedras, Marcer, Alonso-Blanco, & Picó, 2014). In other words, genetically distinct groups are thought to have emerged largely because of dispersal limitation and genetic drift rather than geographically distinct selection pressures. Under an IBD model, we would expect to see a strong association between genetic distance and geographic distance in Hawaii. However, our data deviate from the expectations of IBD in two important ways. First, we see evidence of a single genetic group (orange) spread across nearly all sampling sites with no apparent geographic association. Second, another genetic group (purple) appears at just two sampling sites that are separated by over 500 km, nearly the maximum distance possible across our sampling sites. Dispersal distances for *C. elegans* have not been measured precisely, but numerous lines of evidence suggest they are greater than might be expected for such a small animal. By themselves, nematodes have been observed moving over relatively short distances, *e.g*., one meter in soil over a month period (Barrière & Félix, 2007). However, it was recently shown that wind-mediated transport could facilitate long-range dispersal for nematodes less than 0.75 mm in length, especially under high humidity conditions (Ptatscheck, Gansfort, & Traunspurger, 2018). *C. elegans* dauer larvae are typically 0.4 mm in length, desiccation resistant, and capable of surviving without feeding for months (Cassada & Russell, 1975; Klass & Hirsh, 1976). Furthermore, *C. elegans* has been isolated in association with several terrestrial invertebrate species, including snails, slugs, and isopods, which are thought to act as dispersal vectors (Petersen, Hermann, et al., 2015; Schulenburg & Félix, 2017). Finally, global patterns of genetic diversity in the species implicate human-assisted, long-range dispersal as the likely explanation for many isotypes sampled from locations at least 50 km apart (Lee et al., 2021). The isotype ECA251 could be an extreme case of long-range dispersal as it was isolated from locations that are over 13,000 km apart (California in 1973 and southern Australia in 1983). Moreover, we previously reported evidence of gene flow between Hawaii and the west coast of North America (Crombie et al., 2019). Interestingly, only the widely distributed Hawaiian orange group contains individuals harboring the megabase-scale selective sweeps that dominate European samples. Therefore, we can speculate that an elevated dispersal capacity might explain the near-ubiquitous distribution of the orange group in Hawaii. Considering the geographic distribution of the purple group, we suspect that other factors cause this elevated dispersal. Given the striking similarity in habitat and climate variables at the two sites where the purple group was sampled, we suspect that the unique selection pressures associated with this type of environment might underlie the shared ancestry. In other words, local adaptation to this habitat likely contributes to the patterns of genetic diversity and differentiation we observed. Evidence of local adaptation does not rule out that IBD contributes to the patterns of Hawaiian diversity because local adaptation and IBD are not mutually exclusive. For example, recent studies of *A. thaliana* show that IBD and local adaptation each contribute to patterns of genetic differentiation at the regional scale on the Iberian peninsula (Castilla et al., 2020).

### The genomic architecture of local adaptation

Population structure and complex demographic histories can mislead efforts to identify signatures of local adaptation. However, methods to make GEA more robust to various types of structure exist. In the case of a GEA using GWA, the NemaScan pipeline accounts for the relatedness and genetic structure among strains with a genomic relationship matrix that can at least partially account for population structure (Widmayer et al., 2021; Yang et al., 2011). On the other hand, BayPass uses Bayesian hierarchical models to explicitly account for the scaled covariance matrix of population allele frequencies (Gautier, 2015). However, in either approach, the inferences made when using these tools are subject to limitations. For example, simulations testing the performance of the NemaScan pipeline with various mapping panels indicate that false discovery rates are higher when strongly differentiated strains are included in the mapping panel rather than more closely related strains (Widmayer et al., 2021). This limitation means that some regions implicated in local adaptation might reflect unresolved population structure among the sample of Hawaiian strains. A similar limitation applies to BayPass, wherein spurious signals of local adaptation to environmental variables could be caused simply by demography or drift rather than local adaptation. Conversely, corrections for structure can also reduce the power and increase the rate of false negatives in GEA, especially when major axes of genetic variation and selection coincide. Simulations indicate that higher rates of selfing causes greater neutral divergence between populations and can reduce the power of statistical methods to detect local adaptation loci (Hodgins & Yeaman, 2019)

Another consideration for GEA studies is that the environmental parameters measured may just be correlated with other selective parameters of the environment. In these cases, the true aspects of the environment imparting selective pressures cannot be known without further experiments. For example, suppose elevation gradients drove differences in microbial communities in Hawaiian habitats, as they do in other regions (Tang et al., 2020). Under this hypothetical scenario, differences in pathogenicity or nutritional quality of microbial communities might be the true driver of variation in selection pressure for Hawaiian *C. elegans* along elevation gradients.

In this study, we found that regions of the genome implicated in local adaptation to environmental parameters on the Hawaiian Islands frequently overlap with hyper-divergent regions (Lee et al., 2021; Thompson et al., 2015). Hyper-divergent regions are significantly enriched for genes that modulate sensory perception and responses to pathogens in wild habitats (Lee et al., 2021). For example, genomic loci overlapping with hyper-divergent regions underlie natural variation in responses to the pathogens *Nematocida parisii* and Orsay virus (Ashe et al., 2013; Balla, Andersen, Kruglyak, & Troemel, 2015). For these reasons, we suspect that the selective pressures exerted by local microbial communities, although unmeasured in this study, might be especially important proximal drivers of local adaptation in *C. elegans*. In order to formally test this hypothesis, the microbial communities present at sampling sites should be preserved and characterized when possible.

Regardless of what factors appear to be driving patterns of local adaptation, candidate variants within putatively adaptive loci need to be identified and experimentally validated. Direct proof that a genetic variant causes a fitness advantage in a local environment can only be obtained experimentally (Rellstab et al., 2015), but these experiments are difficult to do in a laboratory setting. In plants, reciprocal transplant studies have been used to validate local adaptation to climate (Ågren & Schemske, 2012; Postma & Ågren, 2016). However, this design is challenging in *C. elegans* because methods for transplanting and resampling individuals have not been developed. As an alternative, validation experiments can be performed in the laboratory by exposing experimental populations to environmental extremes and recording allele frequencies for particular variants over time. This approach has been used successfully to validate fitness advantages of specific variants under anthelmintic drug exposure (Dilks et al., 2020; Dilks, Koury, Buchanan, & Andersen, 2021; Hahnel et al., 2018) and to climate variables at collection sites (Evans et al., 2017). The disadvantage is that the complexity of the real environment is not recapitulated in the laboratory, and fitness advantages observed might not translate to natural conditions. The advantage is that the genetics and the environment can be exquisitely controlled in the laboratory. In the future, these validation techniques can probe whether candidate loci contain functional genetic variation that contributes to environmental adaptation.

## Acknowledgements

We thank the members of the Andersen lab for editing the manuscript. We are grateful to the landowners who gave us permission to collect nematodes on their property. We would also like to thank the Hawaii Department of Land and Natural Resources as well as the Natural Area Reserves System for permitting, support for these studies, and general advice about the Hawaiian Islands. This research was supported by start-up funds from Weinberg College of Arts and Sciences and the Molecular Biosciences department at Northwestern University.M.A. was supported by a National Science Foundation CAREER Award (MCB-1552101). E.C.A. is supported by a National Science Foundation CAREER Award (IOS-1751035). The *C. elegans* Natural Diversity Resource is supported by a National Science Foundation Living Collections Award to R.E.T. and E.C.A. (1930382).

## Data accessibility

All the data and code used to perform our analyses and figures are available for download at https://github.com/AndersenLab/molecular_ecology_manuscript with the exception of NemaScan code, which is available at https://github.com/AndersenLab/NemaScan. The variant sets used in our analyses are subset from the variant set released on 20210121 and available at https://www.elegansvariation.org/data/release/latest.

## Author Contributions

T.A.C, E.C.A designed the research. T.A.C, R.E.T, K.S.E, C.M.B, D.E.C, C.M.D, L.A.S, D.L, S.Z, G.Z, N.M.R, M.A., and E.C.A. performed the research. T.A.C, P.B, and K.S.E analyzed the data. T.A.C. and E.C.A. wrote and edited the paper.

## Supplemental Materials

**Supplemental Table 1.** Distinct collection type counts by project. Each collection class is shown by row and each collection project is shown by column. Counts in each field represent distinct substrates for which a collection class was isolated. In many cases, more than one collection type was isolated from a single substrate so the totals are greater than the actual number of substrates collected (4,506).

|  | 2017-Aug | 2018-Oct | 2019-Aug | 2019-Oct | 2019-Dec | 2020-Jan | Total |
| --- | --- | --- | --- | --- | --- | --- | --- |
| *C. elegans* | 38 | 42 | 1 | 6 | 75 | 0 | 162 |
| *C. briggsae* | 95 | 11 | 2 | 10 | 44 | 2 | 164 |
| *C. oiwi* | 12 | 5 | 0 | 0 | 14 | 0 | 31 |
| *C. tropicalis* | 13 | 5 | 0 | 4 | 4 | 0 | 26 |
| *C. spp.* | 0 | 0 | 0 | 0 | 13 | 0 | 13 |
| *C. kamaaina* | 2 | 0 | 0 | 2 | 0 | 0 | 4 |
| non-*Caenorhabditis* | 724 | 208 | 0 | 118 | 318 | 3 | 1371 |
| unknown nematode | 958 | 340 | 75 | 310 | 634 | 7 | 2324 |
| no nematode | 693 | 128 | 19 | 130 | 230 | 7 | 1207 |
| **Total** | 2535 | 739 | 97 | 580 | 1332 | 19 | 5302 |

**Supplemental Figure 1.**
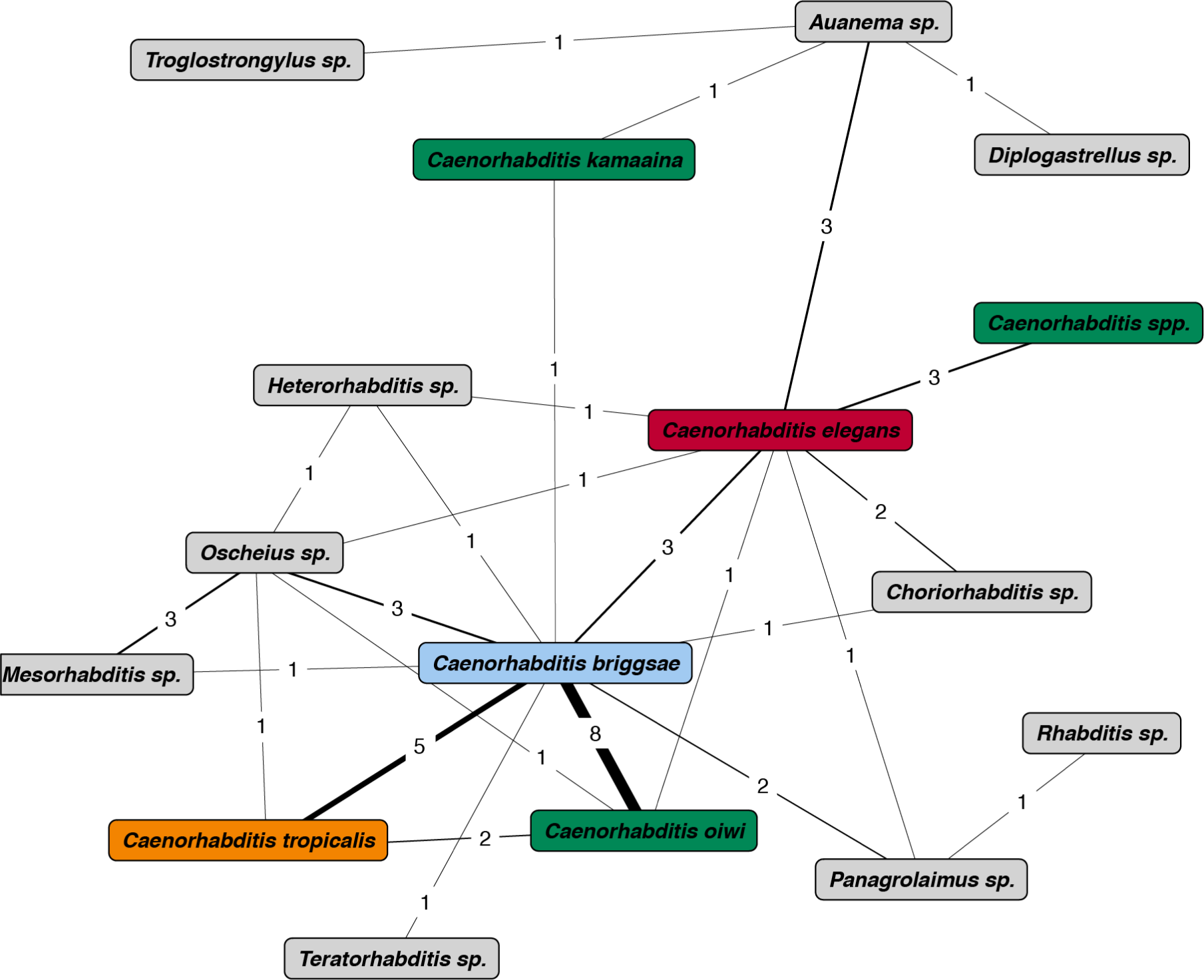
Cohabitation network of nematode taxa. Cohabitation network with taxa as nodes. Node colors represent taxa (red is *C. elegans*, blue is *C. briggsae*, orange is *C. tropicalis*, other *Caenorhabditis* species are green, non-*Caenorhabditis* species are grey). The edges connecting nodes indicate the number of cohabitations between the nodes. The edge width corresponds to the number of cohabitations.

**Supplemental Figure 2.**
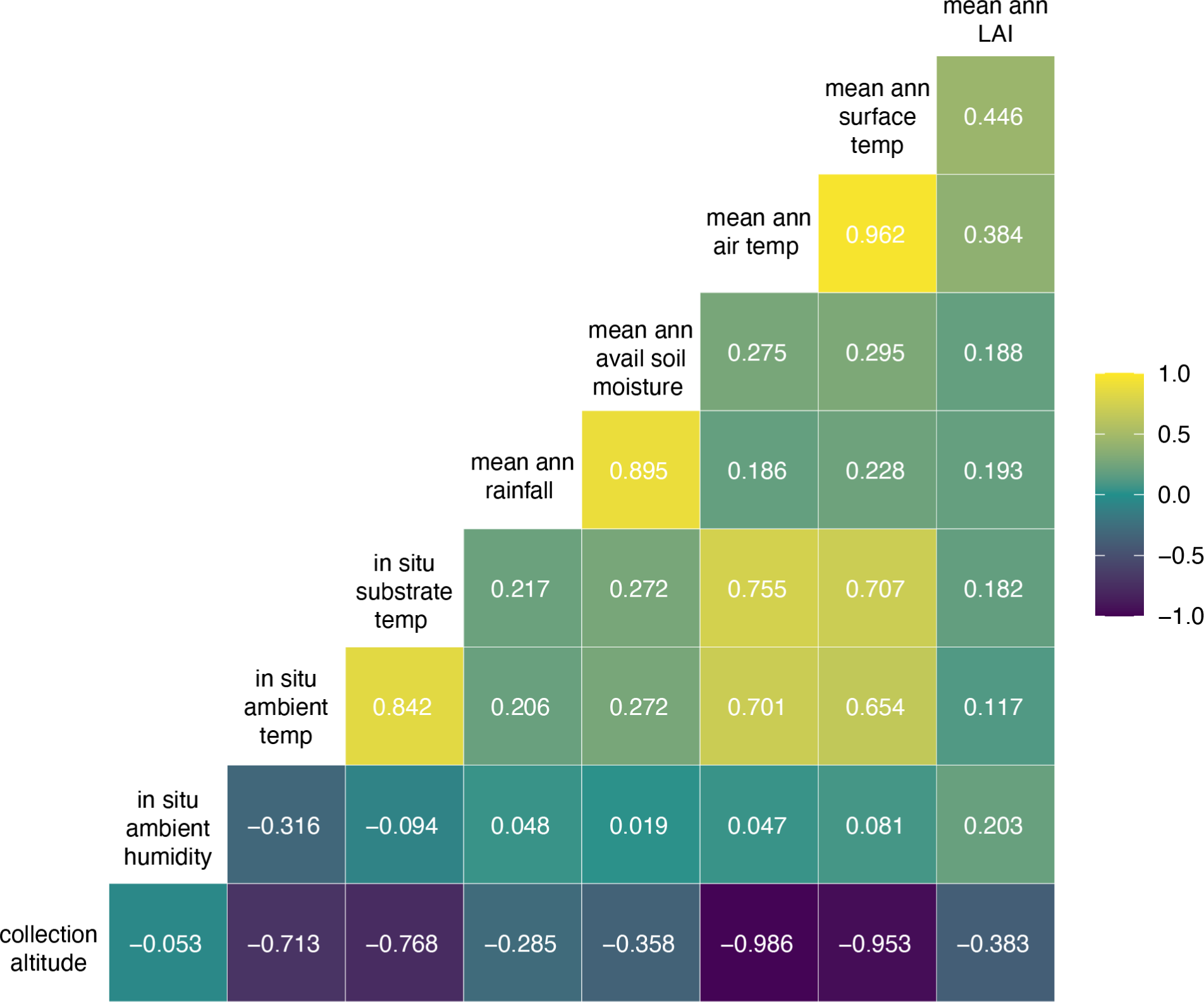
Environmental parameter correlations. A correlation matrix for the continuous environmental parameters is shown. The parameter labels for the matrix are printed on the diagonal, and the Pearson correlation coefficients are printed in the cells. The color scale also indicates the strength and sign of the correlations shown in the matrix.

**Supplemental Figure 3.**
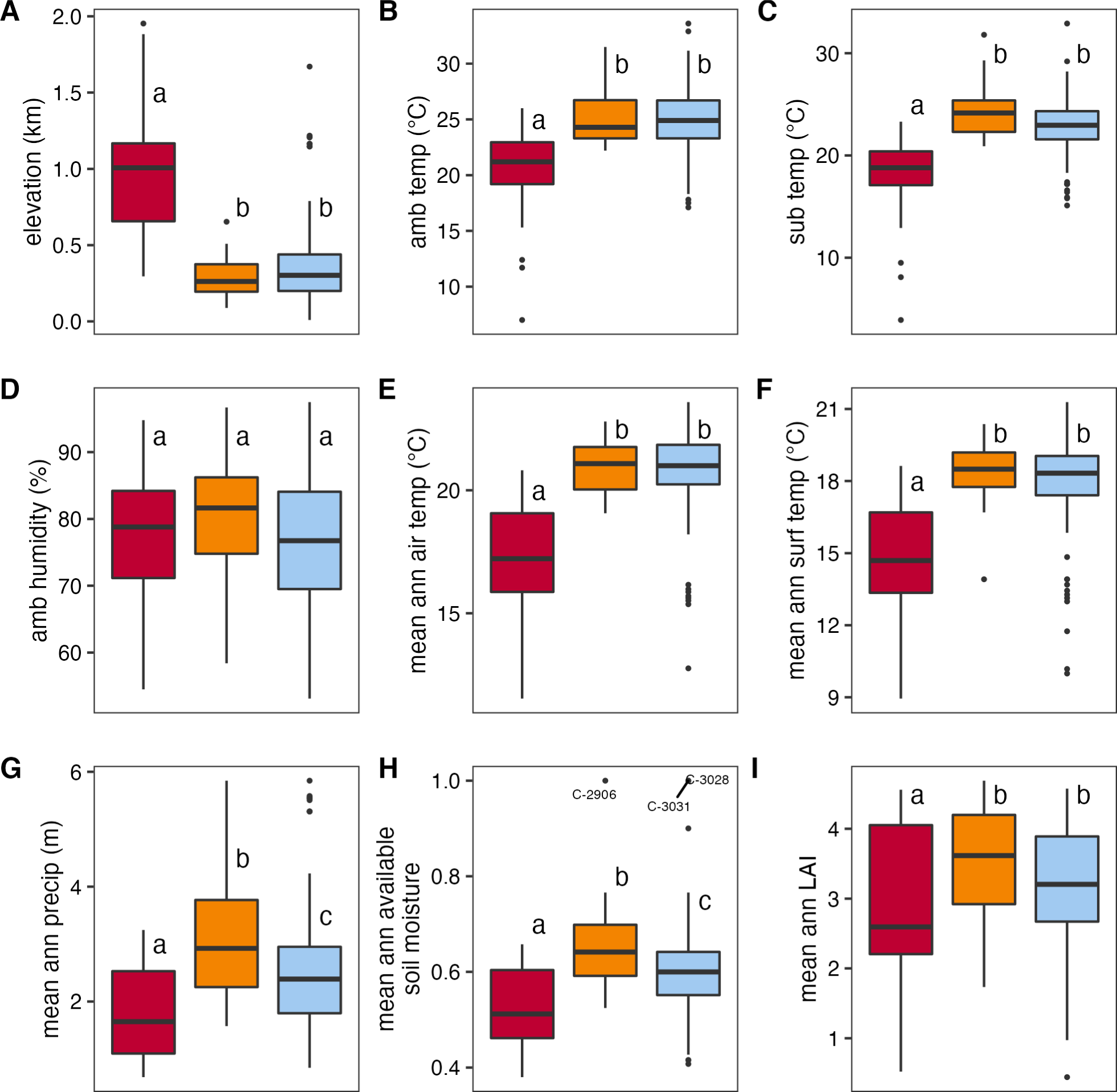
Environmental parameters for selfing *Caenorhabditis* collections. Environmental parameter values measured at the time of collection: elevation (A), ambient temperature (B), substrate temperature (C), and ambient humidity (D). Environmental parameter values obtained from environmental models: mean annual air temperature (E), mean annual surface temperature (F), mean annual precipitation (G), mean annual available soil moisture (H), mean annual leaf area index (I). Tukey box plots are plotted by species (red = *C. elegans*, orange = *C. tropicalis*, blue = *C. briggsae*) for each environmental parameter. Letters above the boxes summarize statistical significance of comparisons between the species shown. Species with a different letter are significantly different; species with the same letter are not different. Comparisons were made using a Kruskal-Wallis test and Dunn’s test for multiple comparisons with *p*-values adjusted using the Bonferroni method.

**Supplemental Figure 4.**
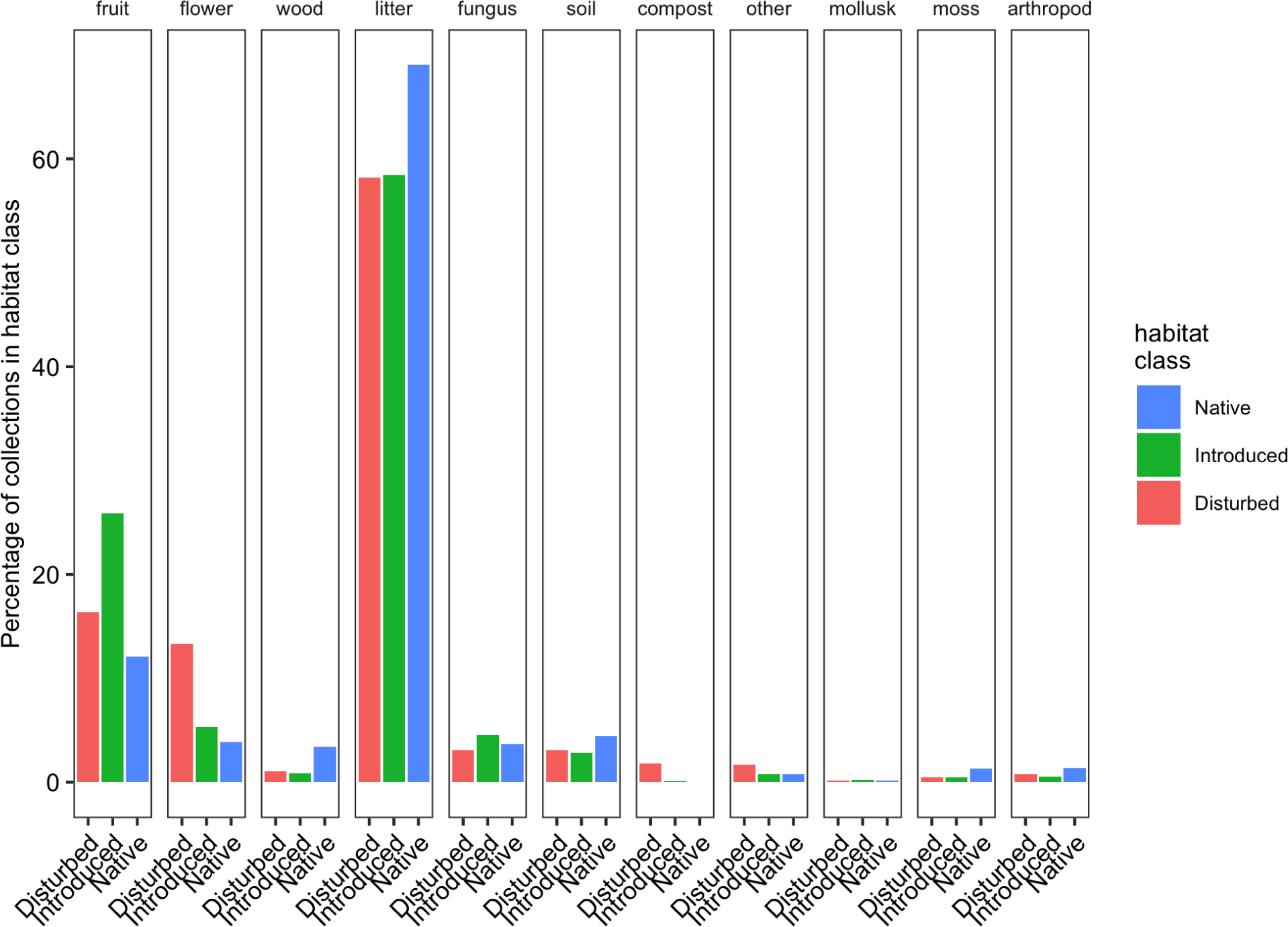
Substrate sampling bias by habitat status. The percentages of collections within each habitat class are plotted by substrate type. Each of the three habitat classes are shown on the x-axis and colored as indicated in the key on the right. Each facet represents a unique substrate type and is labelled on top of the plot. For each habitat class, the bars across all facets sum to 100%.

**Supplemental Figure 5.**
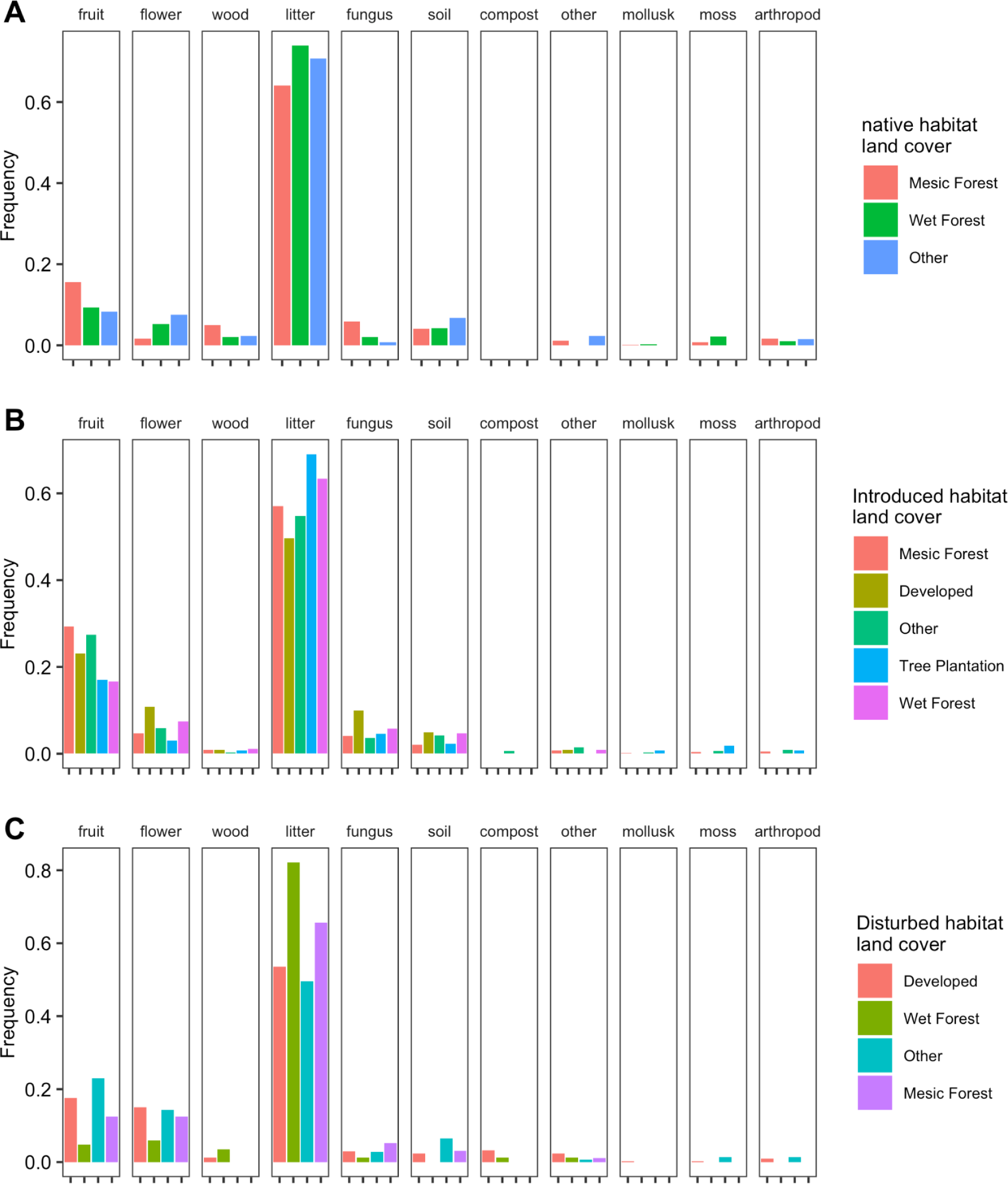
Substrate sampling bias by land cover. The frequencies of collections within each land cover are plotted by habitat class and substrate. The three habitat classes are plotted in (A) native habitat, (B) introduced habitat, and (C) disturbed habitat. Within each habitat class, land cover types are plotted on the x-axis and colored as indicated in the key on the right. The facets represent unique substrate types and are labelled on the top of the plot.

**Supplemental Figure 6.**
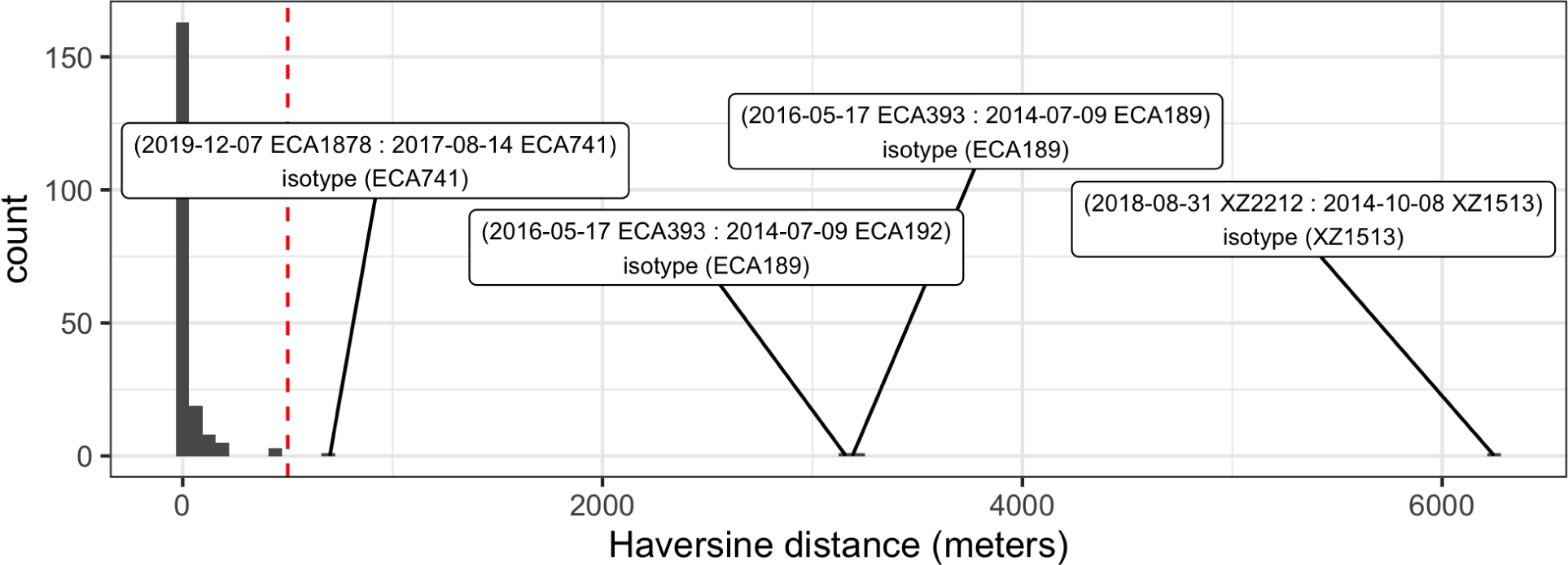
Geographic distance between isotype samples. The histogram shows the counts of pairwise Haversine distances between samples within an isotype. Haversine distance is calculated as the angular distance between two points on the surface of a sphere and is used to account for the curvature of the Earth. Counts are on the y-axis, and physical distance between samples is on the x-axis. The dashed red line indicates 500 meters, only four pairwise distances exceed 500 meters. The collection date and strain name for each pair isolated over 500 meters from each other are shown in the boxes separated by a colon. The isotype for each strain pair is also shown on the second line within the box.

**Supplemental Figure 7.**
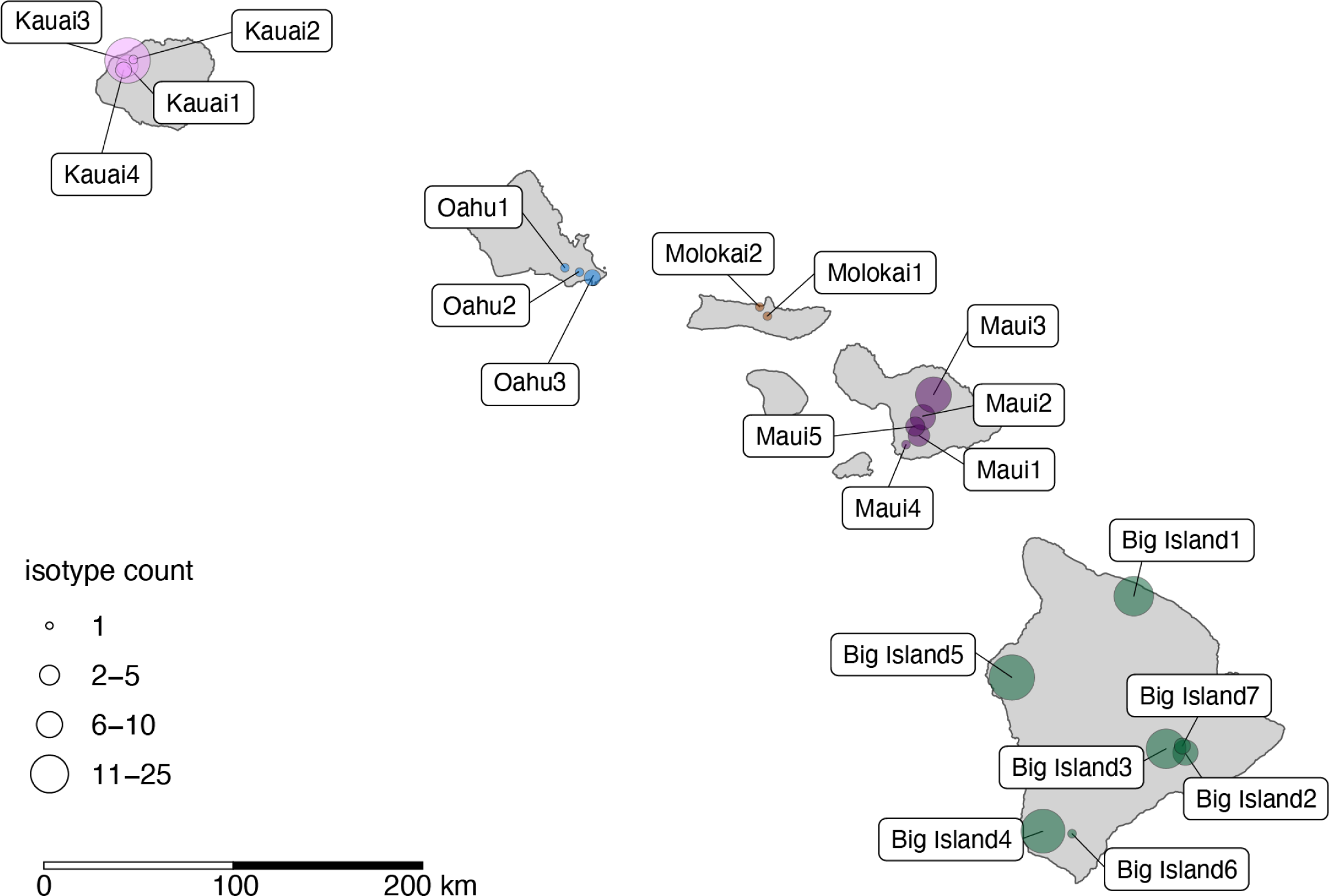
Map of three kilometer sampling locations for *C. elegans*. The 21 three kilometer diameter sampling locations are plotted on a map of Hawaii. The center of each circle indicates the centroid of the sampling location. The size of the circle represents the number of isotypes found within the location. There are seven locations where a single isotype was sampled. The labels indicate the names of each location and are connected to the centroid by a black line. The circles are colored by island.

**Supplemental Figure 8.**
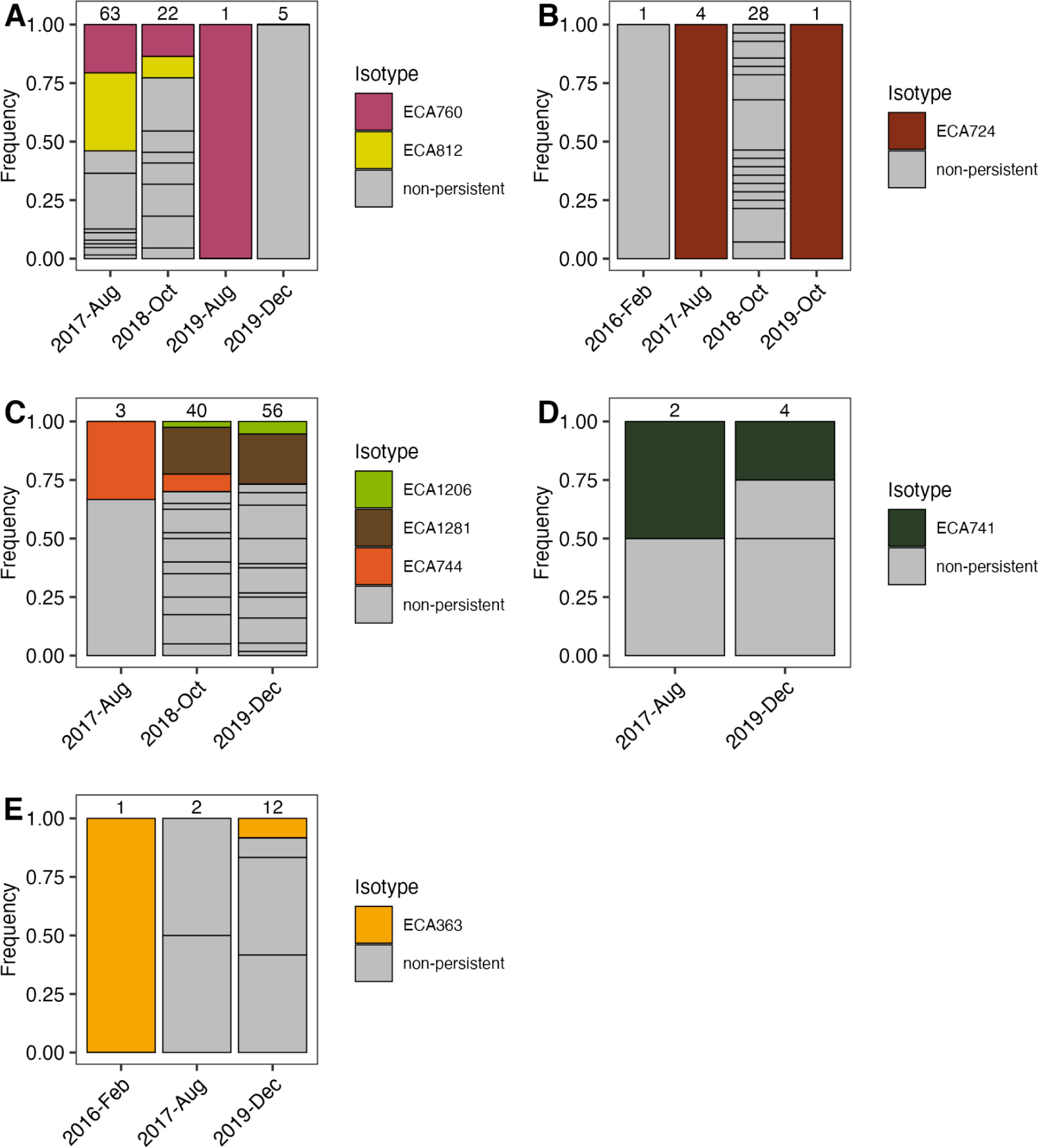
Temporally persistent isotypes by sampling location. Sample frequencies for temporally persistent isotypes are plotted by collection time for each sampling location: (A) Big Island 1, (B) Big Island 3, (C) Big Island 5, (D) Maui 1, (E) Maui 2. Within each sampling location the persistent isotypes are plotted with a unique color corresponding to the key to the right of the plot. Non-persistent isotypes are always plotted in grey. The total number of isotypes sampled for each collection time is shown above the bars. In B and E, the Feb-2016 time point refers to a small sampling effort we did not include as one of the six major sampling efforts.

**Supplemental Figure 9.**
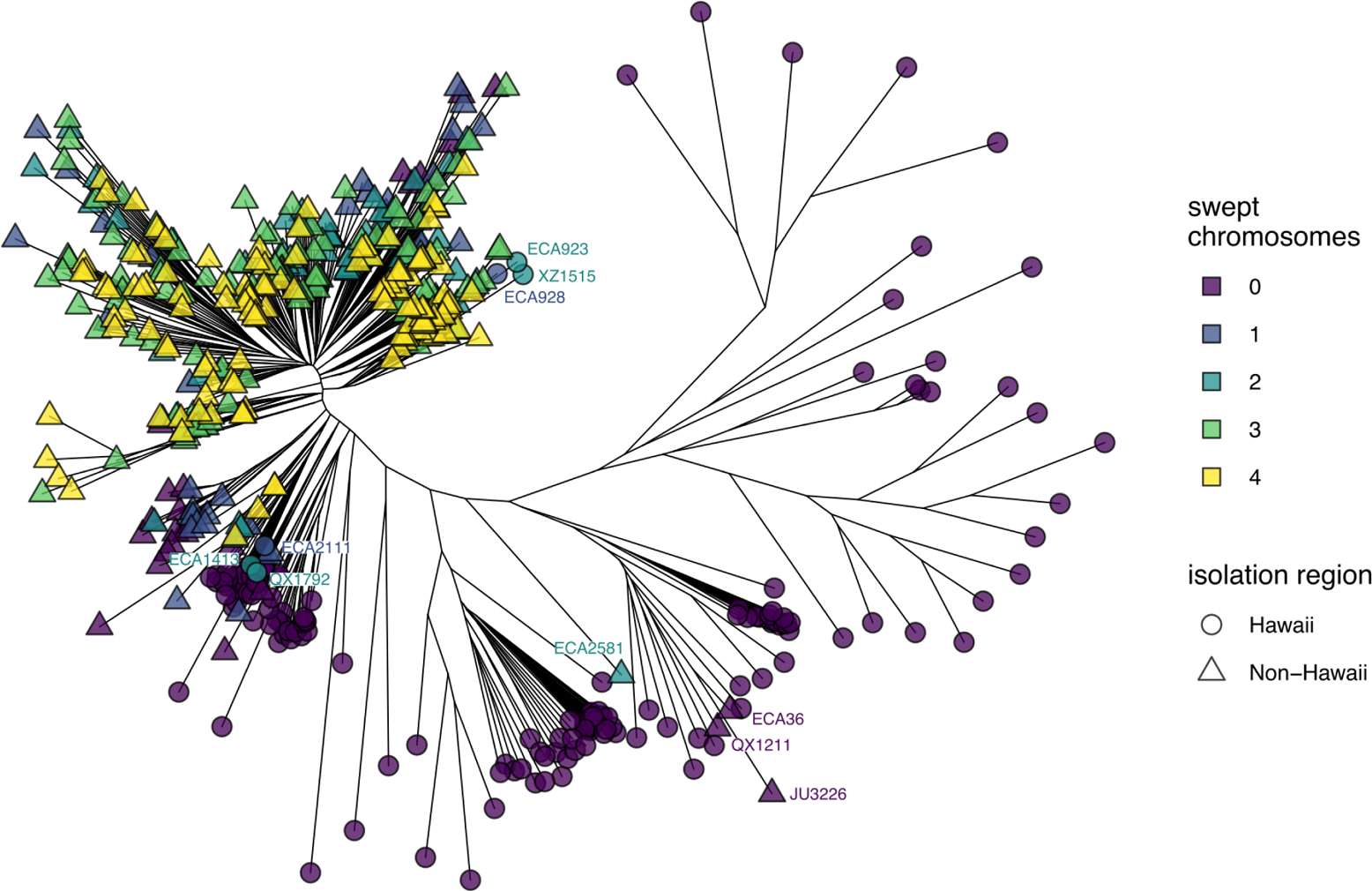
Caenorhabditis elegans genetic relatedness. An unrooted tree built with single nucleotide variants found among the 540 *C. elegans* isotypes sampled from the wild (see Materials and Methods). The branch tips of isotypes sampled from Hawaii are shown with a circle (163), while isotypes sampled from outside of Hawaii are shown with a triangle (377). For each isotype, we classified chromosomes I, IV, V, and X as swept if ≥ 30% of the chromosome consisted of the swept haplotype. Branch tips are colored by the number of swept chromosomes identified for that isotype. The labeled isotypes are either non-Hawaiian isotypes that group with divergent Hawaiian isotypes (ECA2581, ECA36, QX1211, JU3226), or Hawaiian isotypes that contain one or more swept chromosomes (ECA928, ECA923, QX1792, ECA1413, ECA2111, XZ1515). The labels for isotypes are colored by the number of swept chromosomes.

**Supplemental Figure 10.**
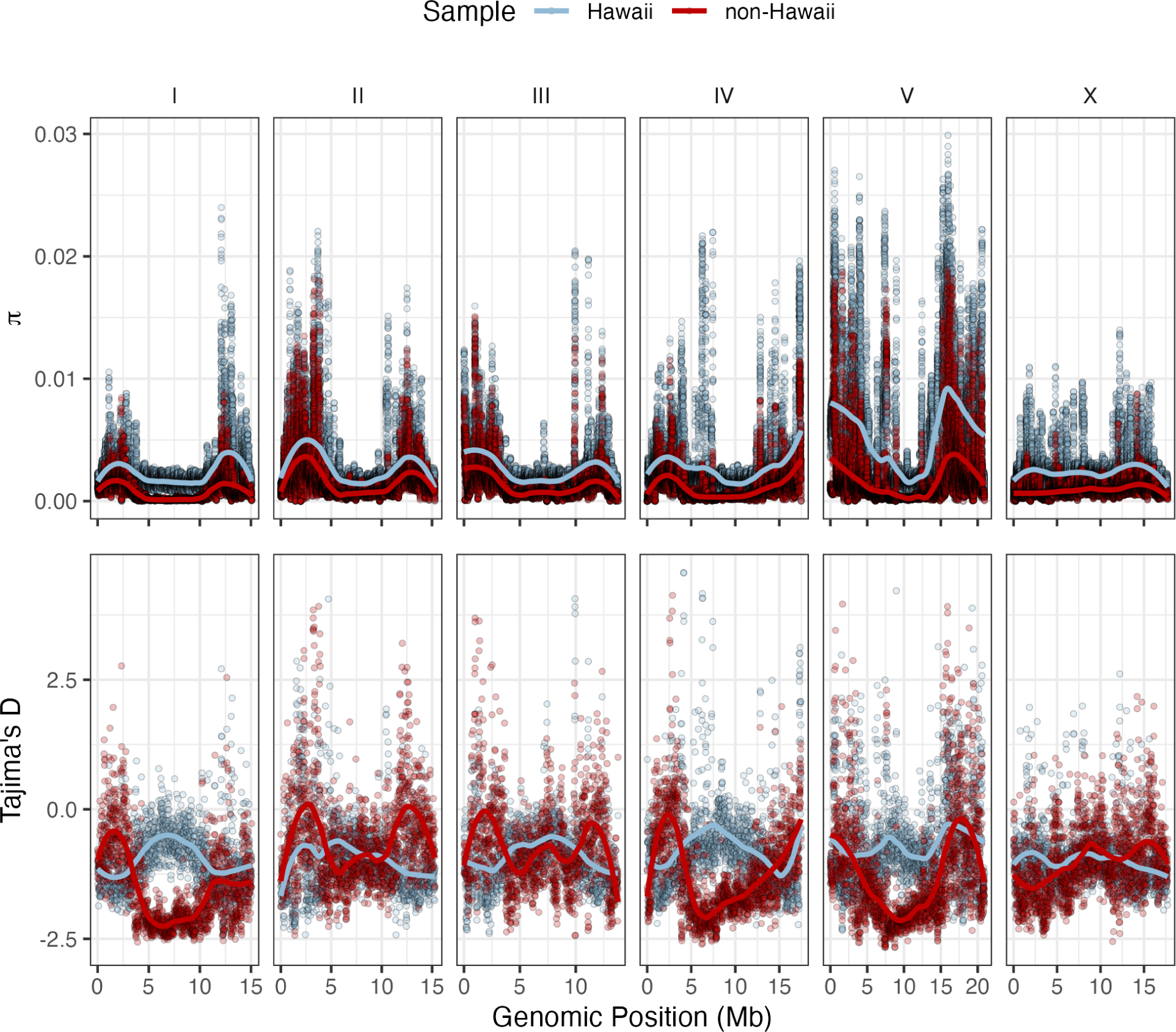
Genome-wide pi and Tajima’s D for Hawaiian and non-Hawaiian samples. (A) Nucleotide diversity (pi) and (B) Tajima’s D measured across the genome for 163 Hawaiian isotypes (blue) and 377 non-Hawaiian isotypes (red) is shown. Genomic position is on the x-axis and facets indicate chromosomes. Nucleotide diversity is measured. Genome-wide pi was calculated along sliding windows with a 10 kb window size and a 1 kb step size. Genome-wide Tajima’s D was calculated with a 10 kb window size and no sliding.

**Supplemental Figure 11.**
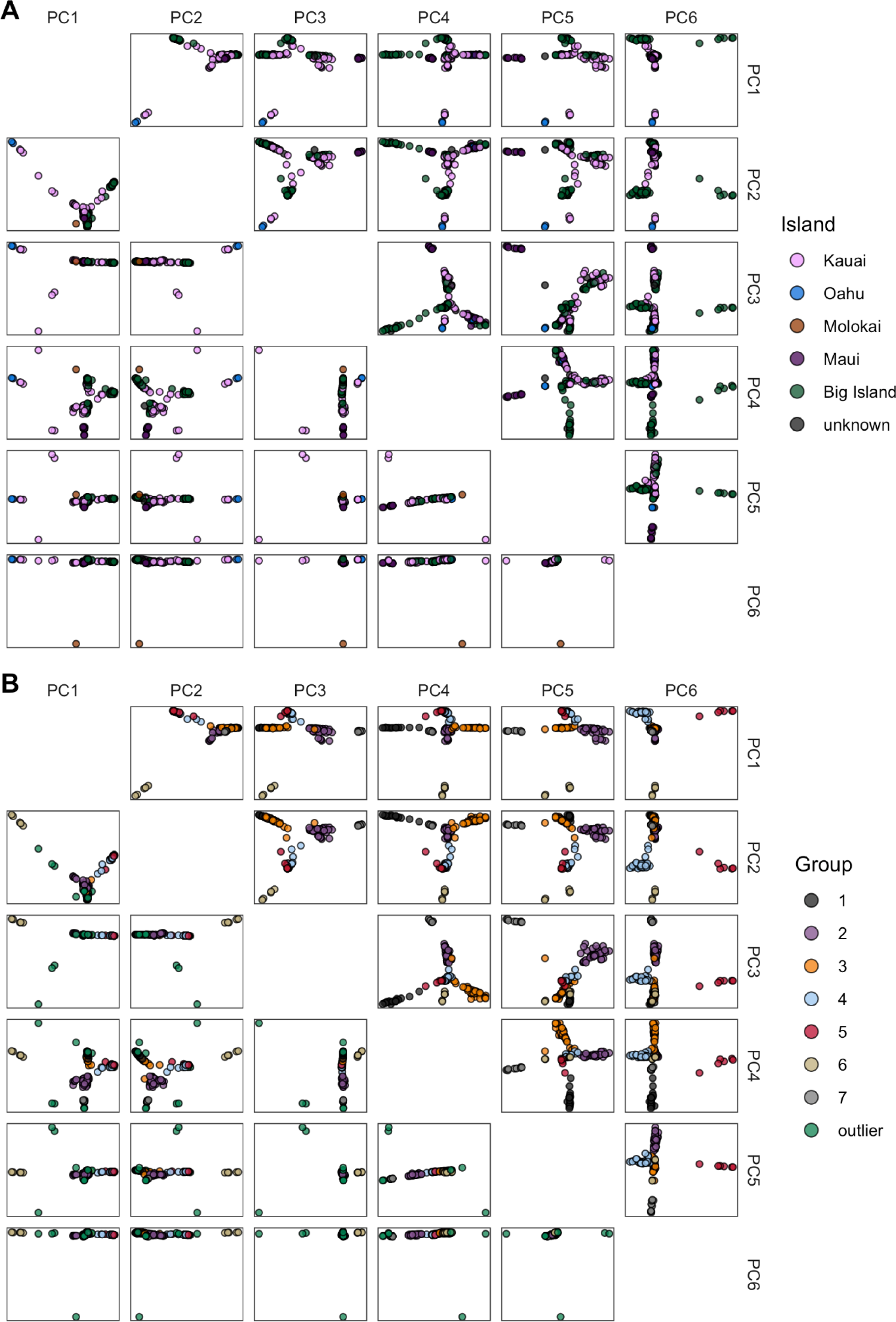
Population structure by principal components analysis. Bi-plots of significant principal components of genetic variation for 163 Hawaiian isotypes. Points represent distinct isotypes, and the axes correspond to values for principal components (PCs) labelled on the top and right of the plots. The plots below the diagonal (open white boxes) show PC values calculated for all 163 isotypes, and the plots above the diagonal show PC values calculated for 149 non-outlier isotypes. **A**) Bi-plots with isotypes colored by island of isolation. **B**) Bi-plots with isotypes colored by genetic clusters revealed by PCA of genetic variation for 149 non-outlier isotypes.

**Supplemental Figure 12.**
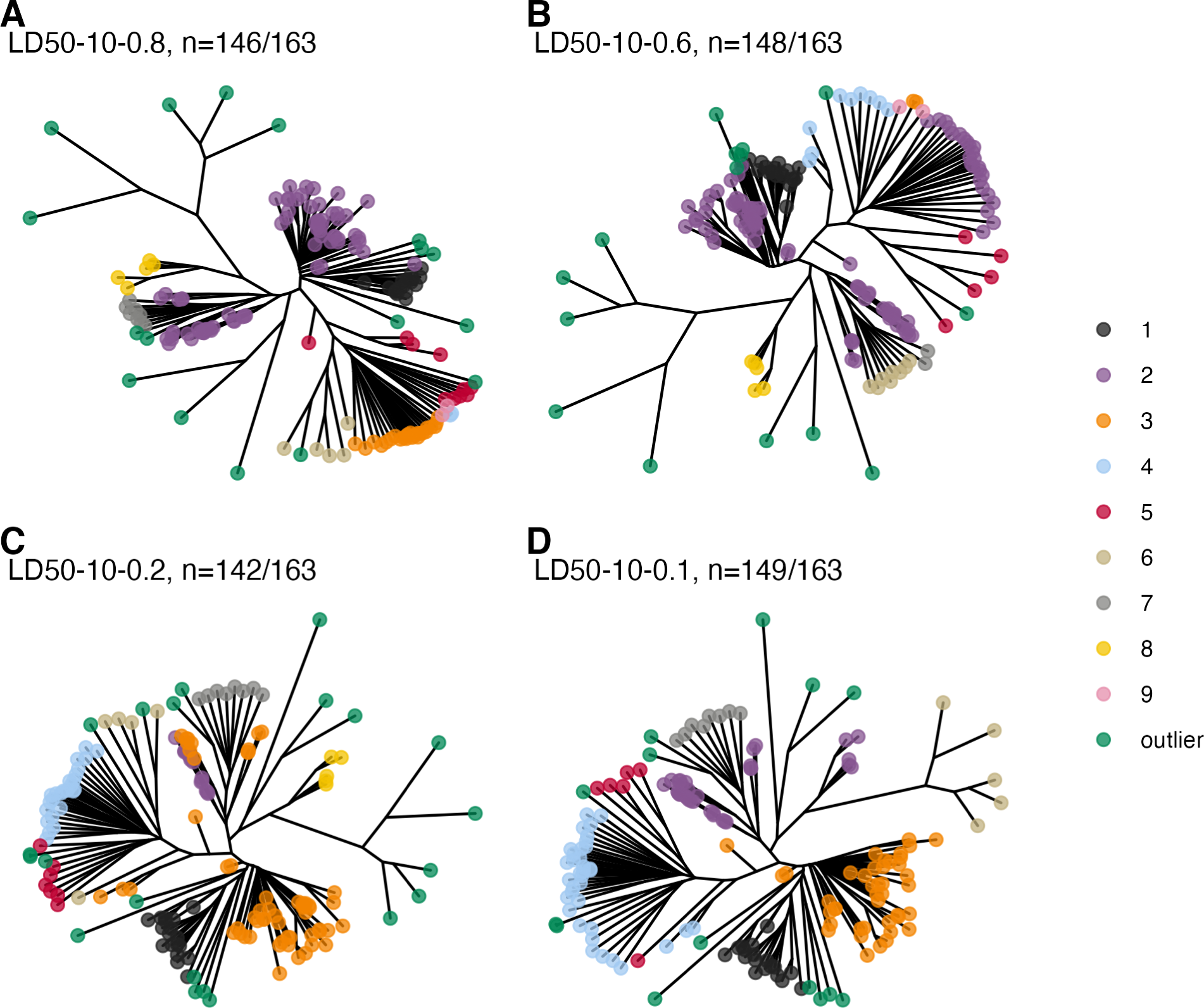
Effect of linkage disequilibrium pruning on population structure inference. **A-D**) Each tree represents the genetic relatedness of the 163 Hawaiian isotypes. The trees are constructed with whole-genome SNV data with varying levels of linkage disequilibrium (LD) pruning thresholds (*r^2^* = 0.8, 0.6, 0.2, and 0.1). Branch tips are colored according to the genetic group to which they are assigned by PCA. The number of isotypes assigned to groups is shown above each plot.

**Supplemental Figure 13.**
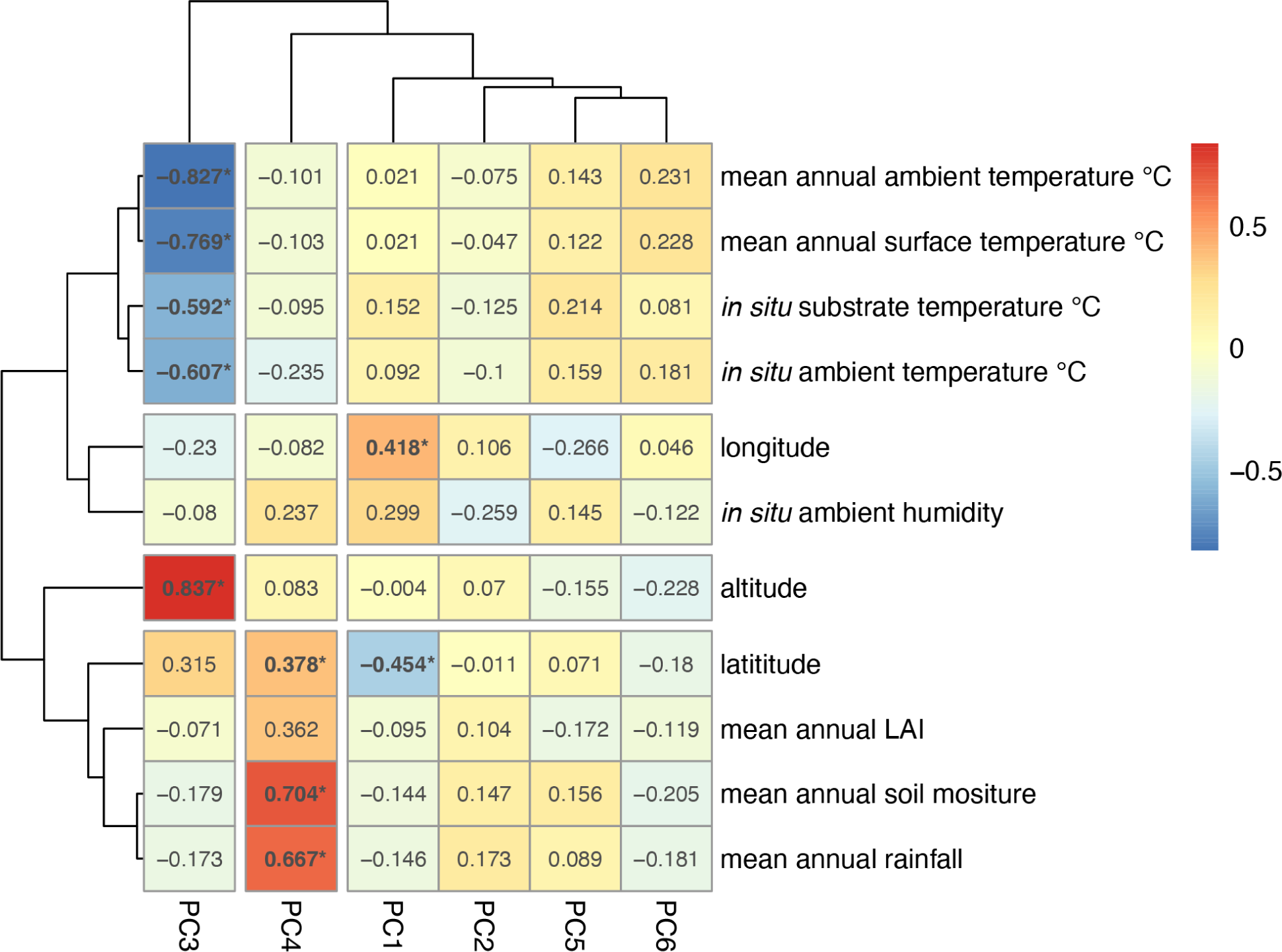
PC and environmental parameter correlation heatmap. The six significant axes of genetic variation (PCs) from PCA on 149 non-outlier Hawaiian isotypes are shown on the x-axis (PCs 1-6), and continuous environmental parameters are plotted on the y-axis. The cell values represent the Pearson’s correlation coefficient between a given PC and environmental parameter values, bold values with an asterisk indicate a significant correlation. The cell colors correspond to the strength and direction of the correlation (see color scale on the right).

**Supplemental Figure 14.**
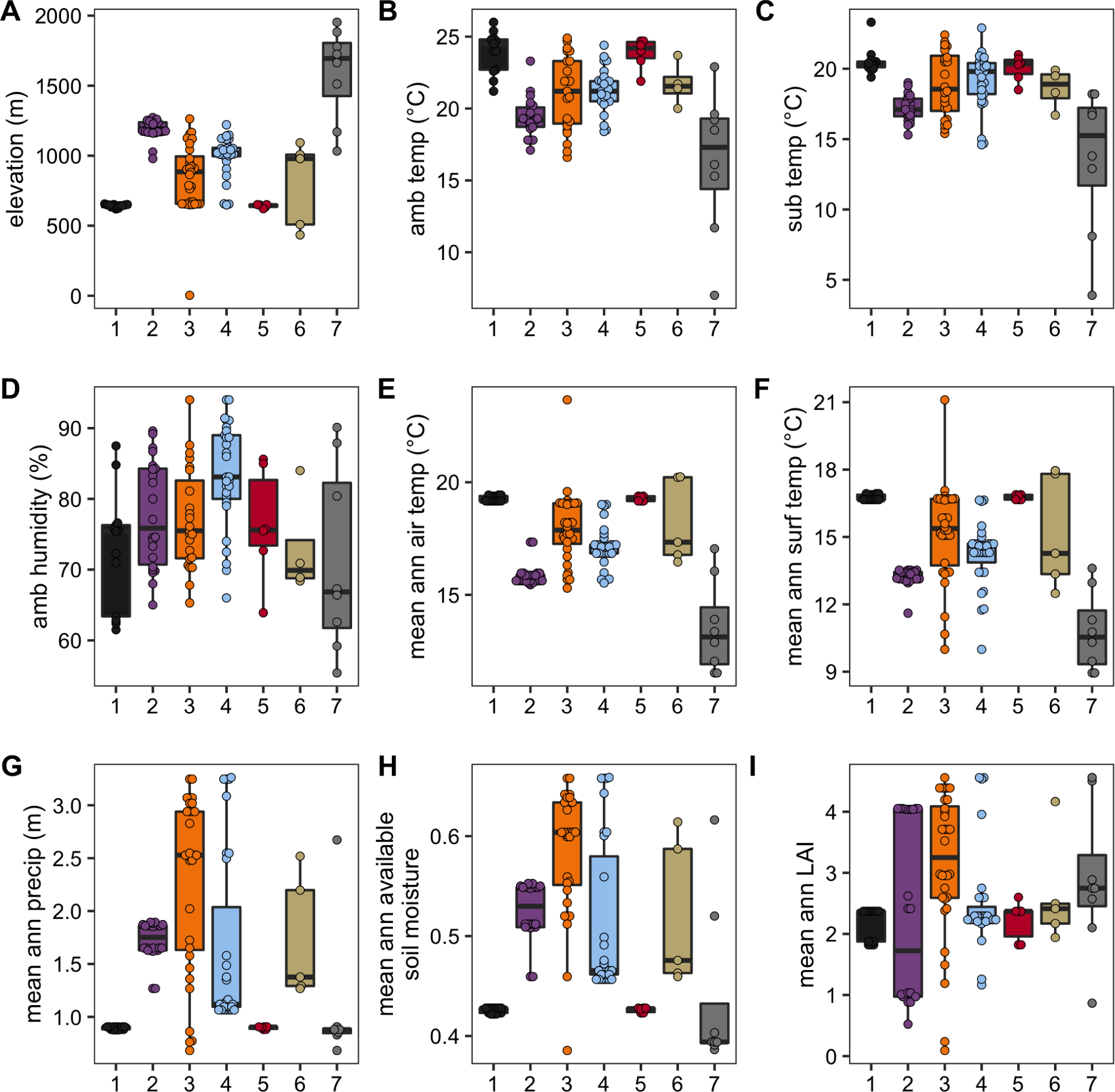
Environmental parameters for *C. elegans* genetic groups. Environmental parameter values measured at the time of collection: elevation (A), ambient temperature (B), substrate temperature (C), ambient humidity (D). Environmental parameter values obtained from environmental models; mean annual air temperature (E), mean annual surface temperature (F), mean annual precipitation (G), mean annual available soil moisture (H), mean annual leaf area index (I). Tukey box plots are plotted by genetic group assignment from PCA (colors) for each environmental parameter.

## Notes

https://github.com/AndersenLab/molecular_ecology_manuscript

### Competing Interest Statement

The authors have declared no competing interest.

